# Large Scale and Stable Graph Differential Analysis via Multi-Layer Node Embeddings and Ranking

**DOI:** 10.1101/2024.06.16.599201

**Authors:** Panagiotis Mandros, Ian Gallagher, Viola Fanfani, Chen Chen, Jonas Fischer, Anis Ismail, Lauren Hsu, Enakshi Saha, Derrick K. DeConti, John Quackenbush

**Author notes:** now at University of Melbourne, Melbourne, Australia (Ian Gallagher). now at Max Planck Institute for Informatics, Saarbrücken, Germany (Jonas Fischer). now at University of South Carolina, Columbia, SC, USA.

## Abstract

Advances in computational biology now enable the inference of increasingly accurate, genome-wide molecular interaction networks from multi-omic data collected across large cohorts. Comparing such networks across distinct biological states can identify biologically informative and potentially actionable higher-order interactions that differentiate these states. However, developing methods for effective graph differential analysis remains challenging as it requires capturing subtle structural changes within the context of complex, high-dimensional networks shaped by heterogeneous biological processes. We introduce node2vec2rank (n2v2r), a multi-layer spectral embedding algorithm that, in contrast to conventional feature-based approaches, compares graphs by inferring node representations that summarize network structure in a data-driven manner. Node2vec2rank is computationally efficient, stable, and provably recovers the correct ranking of differences between weighted graphs. We used n2v2r to compare networks from breast cancer subtypes, to analyze networks capturing single-cell dynamics, and to investigate sex differences in lung adenocarcinoma. The results of these analyses show that n2v2r is a versatile and powerful tool that can uncover biologically relevant insights from complex biological networks, distinguishing between states of health and disease.

## 2 Introduction

Advances in sequencing technologies provide multifactorial data on biological systems, and methods in computational biology can infer graph representations of the associated complex molecular interactions, including those involving gene regulation and epigenetics (Langfelder and Horvath, 2008; Glass et al., 2013; Shutta et al., 2022; Morabito et al., 2023; Saha et al., 2024b). These biological networks can be modeled by integrating multiple complementary data types to create increasingly accurate and fine-grained descriptions of complex biological processes (Silverbush et al., 2019; Schulte-Sasse et al., 2021). Their analysis has provided new insights into complex disease processes (Lopes-Ramos et al., 2018; Reel et al., 2021; Ruiz et al., 2021).

Differential feature analysis remains the primary approach for studying disease. It compares measurements, such as gene expression, between populations representing distinct biological states (for example, tumor versus adjacent normal tissue) to identify features that differ significantly and provide insight into the molecular changes underlying disease. Widely used analytical pipelines rank genes by their ability to distinguish states and include generalized linear model tools such as edgeR (Robinson et al., 2010), DESeq2 (Love et al., 2014), and limma (Ritchie et al., 2015). The resulting ranked lists are typically compared against databases of biological pathways or processes to identify functions that are overrepresented by features at the top of the list and so may define the states being studied (Ashburner et al., 2000; Kanehisa and Goto, 2000; Subramanian et al., 2005).

Adapting this paradigm for graph data and the rich representational power graphs capture, can help identify differences in higher-order molecular interactions, including genes that, although not showing differential expression, may be differently regulated and so drive functional differences between biological states (Lopes-Ramos et al., 2020; Weighill et al., 2021; Saha et al., 2024a). These analyses often infer a network for each biological context and then rank nodes based on differences in their summary statistics. A common choice for such a statistic is the node degree, corresponding to the sum of all edge weights associated with a node, which serves as an interpretable proxy for differences in node connectivity. This “bag-of-features” approach, however, does not fully leverage higher-order graph structures, such as communities and cascades. For example, the degree fails to account for situations in which nodes change connectivity but maintain their overall degree, as can occur when transcription factors alter their relative influence on target genes while preserving their overall activity, or when genes change their correlation partners as cells modify their biological functions.

Representation learning provides a more refined approach to graph comparison than traditional bag-of-feature approaches. Rather than relying on user-defined summary statistics, these methods automatically learn informative features directly from data, aiming to capture the most relevant structural, functional, or semantic patterns in an efficient and expressive form. In the context of graph comparison, representation learning can infer node representations and relevant context-dependent summaries that capture higher-order graph structures in a data-driven manner. Representation-based methods, such as graph statistical modeling (Hoff et al., 2002; Rubin-Delanchy et al., 2022; Arroyo et al., 2021; Gallagher et al., 2021; Levin et al., 2019) and graph machine learning (Grover and Leskovec, 2016; Hamilton et al., 2017; Perozzi et al., 2014; Veličković et al., 2017), often exhibit better performance than feature-based methods in graph tasks, such as node clustering or classification, and allow prediction without having to explicitly choose the node statistics used in modeling.

Not surprisingly, the application of node representation learning to perform differential analysis of network graphs in biological systems presents numerous challenges. When comparing multiple graphs, the problem is often framed as multi-layer graph analysis, requiring the construction of a shared latent space in which nodes from different graphs (or graph layers) can be meaningfully compared (Kivelä et al., 2014). For a single graph, network communities can be defined by affinity and structural equivalences, which translate to node proximities in the latent space. However, there is no consensus on the optimal method for multi-graph comparison to find differences, and the corresponding properties in the latent vector space remain poorly defined. One additional feature—and challenge—of graphs is that they are known to exhibit a no-free-lunch behavior where many “truths” can be conveyed simultaneously (Peel et al., 2017; Priebe et al., 2019), suggesting that multiple meaningful differences may be detectable and should be accounted for in graph comparison.

Lastly, the overall procedure of graph comparison should be computationally efficient, trustworthy, and amenable to easy integration with established computational pipelines. For example, embedding algorithms based on neural networks or random walks are unstable when generating representations (Schumacher et al., 2020; Wang et al., 2022), and little is known about their theoretical properties, such as convergence. Computationally efficient procedures not only facilitate the analysis of multiple large graphs but also support the creation of ensembles consistent with the veridical data science principles of predictability, computability, and stability (PCS) (Yu and Kumbier, 2020).

Our solution to the representation-based graph differential analysis problem is node2vec2rank (n2v2r), a computationally efficient, stable, and theoretically sound method designed to uncover differences in graphs that leverages the rich representation power of graphs. Node2vec2rank uses multi-layer spectral embedding to project the graphs to be compared into a joint latent space where nodes can be ranked based on their representation disparities. This approach combines the clarity of simpler feature-based comparison methods that contrast node summary statistics with the power of data-driven node representations and vector arithmetic to capture higher-order graph structures. As its core embedding algorithm, n2v2r extends the unfolded adjacency spectral embedding (UASE) (Gallagher et al., 2021), a method based on singular value decomposition that efficiently handles multiple large graphs while enabling theoretical guarantees for correctly ranking differences. To ensure stability, n2v2r aggregates rankings derived from varying the number of embedding dimensions and using different distance metrics to measure node disparities. The ranked output is then readily available for downstream applications, such as gene set enrichment analysis and visualization of the most significant differences.

In this manuscript, we describe and validate several important properties of n2v2r. In particular, we formally prove identifiability for the correct ranking of node differences in weighted graphs and two distance metrics, we demonstrate the performance with both single-cell and bulk real-world data and using multiple network inference methods while exploring different biological questions and applications, and show how n2v2r can be integrated with various downstream analysis tools. The Python code and data analysis notebooks that walk through the analyses presented here are available at https://github.com/pmandros/n2v2r.

## 3 Results

### 3.1 The Node2vec2rank Framework

We are interested in the problem of ranking nodes in unipartite graphs based on how much their “behavior” changes between graphs that represent different biological states. Given a set of *K* graphs with a common set of *n* nodes, we denote their weighted adjacency matrices as **A**^(*k*)^ ∈ ℝ^*n×n*^, such that 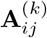 is the weight of edge (*i, j*) in graph *k* ∈ [*K*].

In this formulation, the (weighted) node degree statistic, also known as connectivity, is defined as

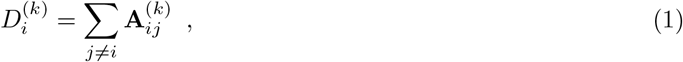

which for a pair of graphs **A**^(1)^ and **A**^(2)^ leads to the Degree Difference (DeDi) ranking

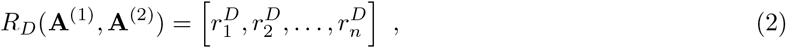

with 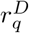 being the node *i* with the *q*-th largest absolute degree difference 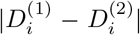 between the two graphs. The degree, given its interpretability, is a widely used node summary statistic for graph comparison; however, it is unable to detect the more complex differences that occur in higher-order graph structures, such as communities and cascades. Consider a scenario in which network graphs are created for two biological states and where some nodes alter their connectivity between graphs while maintaining their overall degree. In this case, comparing node degrees would not identify these differences, but the shift in node connectivity could signify meaningful changes between biological states. In Figure 2, we simulated such a scenario.

#### 3.1.1 Differential Ranking Based on Multi-Layer Node Embeddings

Unlike bag-of-features approaches, which rely on user-defined summary statistics, node embedding algorithms learn high-dimensional node representations that capture higher-order graph structures in a data-driven manner. For a single graph, an embedding function maps nodes to a point cloud in a latent space of *d* dimensions, such that b(**A**) = **X** ∈ ℝ^*n×d*^ where the embedding of node *i* is the vector corresponding to the row **X**_*i*_. We perform graph differential analysis by ranking nodes based on the disparity of their representations between graphs. We refer to this framework as node2vec2rank (n2v2r).

A naive implementation of n2v2r for two graphs would involve creating two node embeddings from separate applications of the embedding function, arriving at

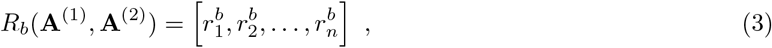

where 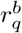 is the node *i* that has the *q*-th largest disparity 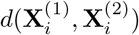 and *d* is some distance metric. While this formulation is straightforward, there is one crucial requirement for n2v2r to be meaningful, which is the construction of a latent space that is provably common for all graphs—otherwise, any comparison would be misleading. For example, applying a naive n2v2r to two gene regulatory networks would generate separate embedding spaces that are not guaranteed to capture the same underlying biology: the first latent dimension of one network might correspond to a metabolic process, while the first latent dimension of the other could reflect cell–cell signaling.

The requirement for a common latent space is satisfied by multi-layer node embedding algorithms b_*M*_ and include statistical models (Gallagher et al., 2021) as well as fine-tuning and transfer learning techniques (Grover and Leskovec, 2016; Dingwall and Potts, 2018). Given a set of graphs 𝒜 = {**A**^(1)^, …, **A**^(*K*)^} with *n* nodes each, these methods create a joint embedding space with *K* sets of *n d*-dimensional represen-tations,

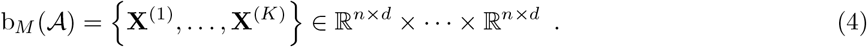

Given two graphs, *A* = {**A**^(1)^, **A**^(2)^}, n2v2r ranks nodes using

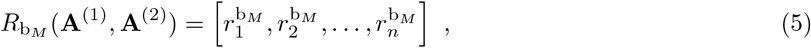

where 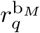 is the node *i* that has the *q*-th largest disparity 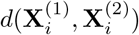 in the joint embedding space generated by b_*M*_.

#### 3.1.2 Ranking Given Multiple Graphs

The ranking process can be naturally extended using vector arithmetic to account for applications with *K >* 2 graphs, a situation that can occur when comparing multiple conditions or tracking the evolution of a series of ordered graphs. In these instances, n2v2r first applies its multi-layer embedding algorithm and then uses three comparison strategies to produce pairwise rankings. The first is a “one-vs-all” strategy in which we perform *K* pairwise comparisons, contrasting the embedding **X**^(*k*)^ with the mean of the remaining embeddings, such that the ranking of node *i* between graph *k* ∈ [*K*] and the rest is a function of

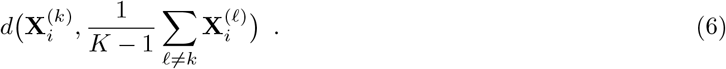

When an ordering for the graphs is available (such as for discrete time points), a second strategy, the “sequential” strategy, compares the embedding **X**^(*k*)^ for *k* ∈ [2, *K*] to the previous embedding **X**^(*k*−1)^. The third comparison strategy, a “one-vs-before strategy,” adopts a stricter notion of ordering (such as for longitudinal data) and compares the embedding **X**^(*k*)^ with the mean of all previous embeddings {**X**^(1)^, …, **X**^(*k*−1)^}

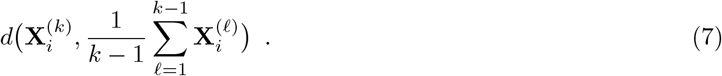

The sequential strategy identifies nodes with the largest transition between consecutive network states. In contrast, the one-vs-before strategy considers all previous graphs and focuses on unique differences for each graph in the ordered set.

#### 3.1.3 Consensus and Stable Ranking

The ranking of nodes in a graph based on their importance remains an open problem. Obtaining a single “true and meaningful” ranking is generally impossible as there is an inherent uncertainty in inferring networks from intrinsically noisy data. Further, even if we assume that a network graph is correct, graphs are inherently complex and can convey multiple truths depending on the graph representation used (Priebe et al., 2019). Node ranking also depends on the choice of embedding dimension and the disparity measure used, both of which can be non-trivial to establish for real-world data. Setting these parameters can be challenging in Life Sciences applications where one wants to balance the predictability, computability, and stability of a method with the need to identify interpretable factors associated with each of the biological states under study (Yu and Kumbier, 2020).

In n2v2r, we address this by generating a collection of rankings using the cross-product of different choices for embedding dimensions and disparity functions. These rankings can then be aggregated into a single stable consensus ranking. As an aggregate function, we use the Borda voting scheme that outputs the mean rank of every node across all rankings. Note that the main requirement for enabling stability given arbitrarily large graphs is computational efficiency, meaning that the individual applications of n2v2r should be tractable. In Figure 1, we illustrate the n2v2r framework for a simulated case-control study with two graphs.

**Figure 1.**
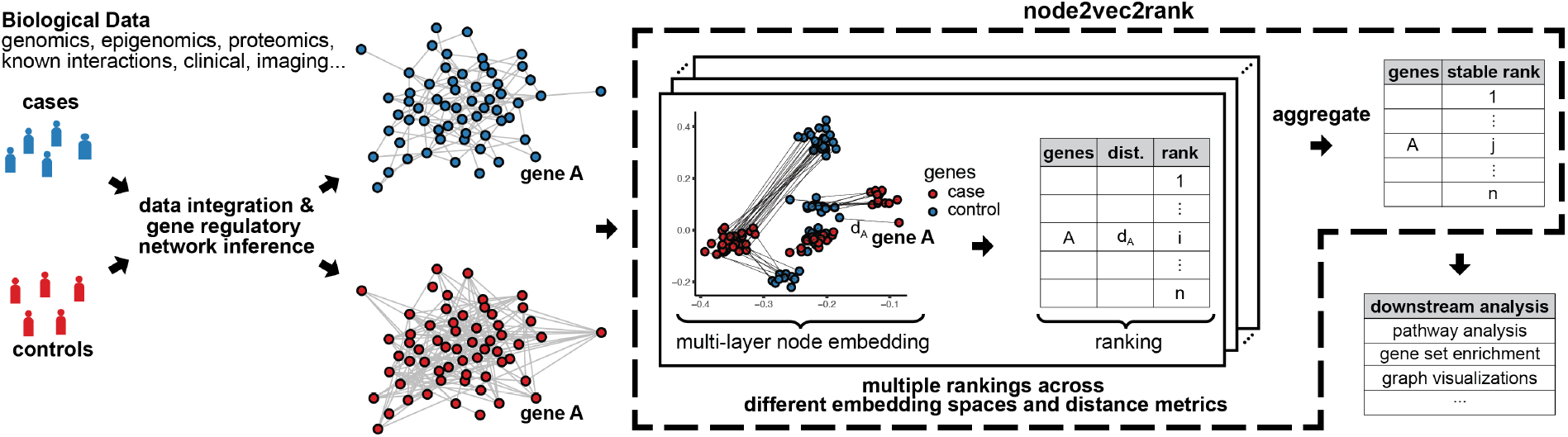
Illustration of node2vec2rank and an analysis pipeline for a case-control study with two graphs. Given biological data, two networks are computationally inferred and subsequently analyzed using n2v2r to identify their differences. Node2vec2rank uses a multi-layer node embedding algorithm to create two sets of vector representations for all genes (depicted here in two dimensions). For every gene, n2v2r computes the disparity between its two representations, which is then used to rank the genes in descending order of disparities. The process is repeated multiple times, producing embedding spaces of varying dimensionalities (for example, 2, 4, 8, 16, …) and rankings based on different distance metrics (cosine, Euclidean, etc.). The multiple rankings are then aggregated to produce a final stable ranking. The output can then be used in downstream applications, including visualizing the largest differences and performing gene set enrichment analysis.

#### 3.1.4 Node2vec2rank with Unfolded Adjacency Spectral Embedding

Reliable and effective tools for analyzing complex scientific “big data” should be scalable and exhibit sound theoretical performance. For graph differential analysis, this means we need guarantees that the method retrieves an optimal ranking under mild assumptions and that the approach is scalable for multiple large graphs and ensembles in a manner that provides stability. To satisfy these criteria, we extend the unfolded adjacency spectral embedding algorithm (UASE) (Gallagher et al., 2021). UASE is an attractive option in our multi-layer setting as it offers longitudinal stability, meaning that if a node has similar behavior in graphs **A**^(*k*)^ and **A**^(*ℓ*)^, then its node embeddings 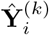 and 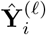 for *i* ∈ [*n*] will be approximately equal. Moreover, SVD is a well-understood, widely used, and scalable algorithm with multiple implementations. Finally, as the UASE in our weighted case is derived from the SVD of the weighted unfolded adjacency matrix representation, it is possible to derive statistical guarantees about the behavior of the resulting embeddings. To this end, we contribute Theorems 1, 2, the proofs of which, together with the necessary model assumptions, can be found in the Methods section.

Theorem 1 states that UASE 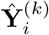 converges to the *d*-dimensional representation of the latent position 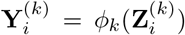 up to an orthogonal transformation, extending previous UASE results for multiple un-weighted binary graphs (Gallagher et al., 2021; Jones and Rubin-Delanchy, 2020) and a single weighted graph (Gallagher et al., 2023).

##### Theorem 1

(Uniform consistency of UASE). *Under Assumptions 1, 2 and 3, there exists an orthogonal matrix* **W** ∈ 𝕆 (*d*) *such that, for all k* ∈ [*K*],

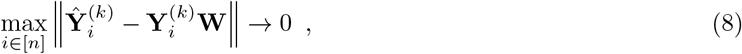

*with high probability as n* → ∞.

In the second part of n2v2r, we rank nodes based on node embedding disparities 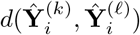 that are estimates of the latent position distances 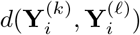 in the embedding spaces. Theorem 1 shows that the latent position distance estimate can only be consistent if the distance function is invariant up to orthogonal transformation, such that for all **W** ∈ 𝕆 (*d*), we have that

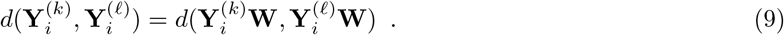

Theorem 2 shows that fitting n2v2r with UASE is consistent for the Euclidean and cosine distances, both of which satisfy the invariance condition.

##### Theorem 2

(Uniform consistency of n2v2r). *Under Assumptions 1, 2 and 3, for all k, ℓ* ∈ [*K*],

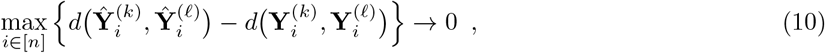

*with high probability as n* → ∞ *for Euclidean distance, and cosine distance provided* 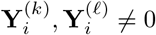.

### 3.2 Analysis of Simulated Networks

We assessed how well n2v2r could detect differences in simulated networks for which we have a known ground truth. While simulated data is hardly an exhaustive testbed for real biological applications, it helps us assess the n2v2r performance in a quantitative manner and provides a framework for exploring where summary statistics fail in differential network analysis.

We explored network differences in two different regimes, termed degree-naive and random. The first corresponds to graph differences where the structure and connections can change, but the degrees of nodes vary little, which is a key motivation for this work as, for many biological situations, much of the network represents essential functions that cannot be disrupted, so the overall structure of the network cannot be altered substantially–although meaningful changes do occur in some connections. In the second, we modeled random differences by allowing the degrees to vary arbitrarily. This represents the type of condition one might expect for a system that is undergoing rapid changes, and so may exhibit changes in network associations that reflect changes in the biological processes active in each condition. For comparison, we considered the degree difference ranking (DeDi), n2v2r with embedding dimension *d* = 1 and Euclidean distance as a baseline (n2v2r e1), as well as the resulting method n2v2r with Borda aggregation (n2v2r b) for dimensionalities *d* = {4, 6, 8, 10, 12, 14, 16, 18, 20, 22, 24} with Euclidean and cosine distances (aggregation over 22 different rankings). Note that the baseline n2v2r e1 can be contrasted with DeDi to compare the performance of a one-dimensional bag-of-features approach against learning a one-dimensional feature.

We simulated the differences by altering the probability matrices in a dynamic stochastic block model such that the community probabilities vary between two graphs (see Methods). Our goal was to assess how well a method can retrieve the nodes of the community with the largest displacement in probability space.

To cover a wide range of ratios, we altered the number of communities in *m* ∈ {4, 6, 8, 10} and the number of nodes in *n* ∈ {50, 100, 500, 1000}, with the task becoming more challenging as the number of nodes decreases or as the number of communities increases. For example, the combination *m* = 6 and *n* = 50 implies that there is an average of 8 nodes per community. We sampled 2000 pairs of graphs for each combination of *m, n*, and graph difference regime and generated violin plots for the mean recall in ranking the nodes of the community that changed the most at the top. The results for *m* = 6 are shown in Figure 2 (results for all *m* and *n* are given in Supplementary Material Figures S1, S2).

**Figure 2.**
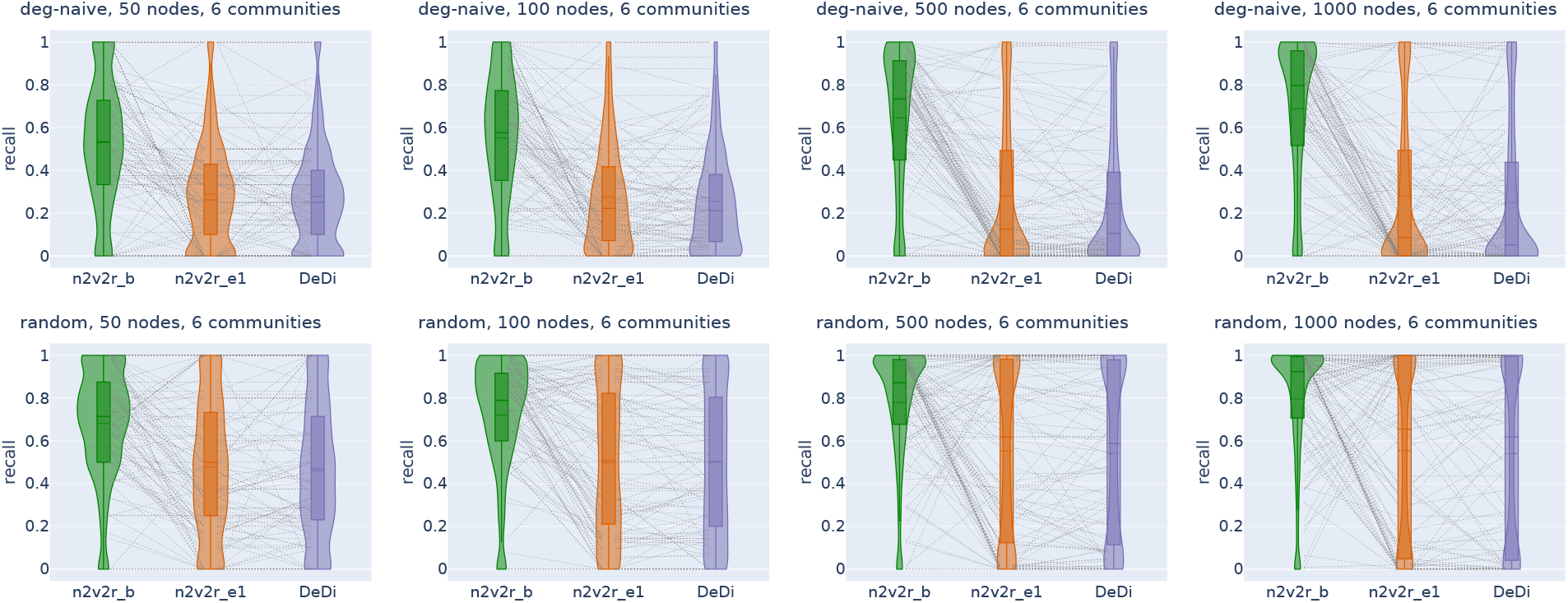
Recall violin plots for retrieving the nodes of the communities with the largest changes between two graphs. The data were simulated using a dynamic stochastic block model with six communities such that the community probabilities vary between the two graphs. The first row corresponds to degree-naive differences, where we permute the probabilities of one community to simulate changing connections, but not node degree. The second row corresponds to random differences, where we randomly change the probabilities of one community. For each scenario, we generated 2000 pairs of graphs by randomly altering the community probabilities as described above. The number of nodes increases along the columns. The goal of this exercise was to retrieve the nodes of the altered community. For a random subset of 100(5%) sampled graphs, we connected the corresponding recall estimates for all methods. The solid and dotted lines in the violins correspond to the median and mean, respectively. The three methods compared are the proposed Borda aggregated n2v2r b, the one-dimensional n2v2r e1 with Euclidean distance, and the degree difference ranking DeDi.

In the first row of Figure 2 with the degree-naive networks we find that, as expected, DeDi is unable to find the target nodes, the one-dimensional n2v2r has slightly better performance, and the aggregated n2v2r performs the best and exhibits the most stable performance, which improves with the number of nodes. For the networks with random differences in the second row of Figure 2, all methods exhibit improved performance, which is unsurprising since the degree-naive regime represents a harder case. We find that the aggregated n2v2r has the best performance, while the one-dimensional DeDi and n2v2r e1 exhibit large variability, with n2v2r e1 performing slightly better. Overall, these results demonstrate that the statistical methods can indeed better adapt to the data, while a single dimension performs poorly at capturing complex graph differences.

We also evaluated performance as a function of dimension, distance metric, and aggregation (Figures S3, S4, S5, S6). We found that both Euclidean and cosine distances perform well, with Euclidean having an advantage because the networks were generated as stochastic block models. Increasing the dimensionality also improved performance in the random differences case, while that is not entirely true for the degree-naive differences, where performance can decline due to overfitting, particularly for a small number of communities.

### 3.3 Analysis of Networks Inferred for Biological Systems

Although analysis of simulated data can provide invaluable insight into the performance characteristics of an algorithm, the true test of any method is whether it can deliver new insights and meaningful results when applied to real biological data. To this end, we considered three distinct applications spanning different network inference methods and data types.

The first study focused on the luminal A breast cancer subtype (lumA BRCA), a disease where metabolic pathways are naturally disrupted during progression. We constructed gene regulatory network models, representing transcription factors (regulators) and their target genes (regulated) as bipartite graphs. Using a unipartite projection, we derived co-regulation networks of genes, which we then used to test n2v2r. In the second example, we used hdWGCNA (Morabito et al., 2023) to analyze cell-cycle data from a single-cell time-course experiment. Rather than using the resulting correlation networks, we focused on hdWGCNA’s topological overlap matrices (TOMs), which estimate similarity between genes based on both correlation strength and shared neighbors. Using n2v2r and DeDi, we examined sequential TOMs to identify network features that changed over time. Lastly, we used data from lung adenocarcinoma to analyze correlation-based networks and to compare these between males and females with the goal of identifying sex-biased groups of genes that might shed light on clinical differences.

Together, these three applications encompass distinct network inference methods and biological contexts—spanning disease subtype comparison, temporal network dynamics, and sex-specific network variation—each posing unique analytical challenges for graph differential analysis.

#### 3.3.1 Characterizing Luminal A Breast Cancer Subtype Mechanisms

In our first system, we used data from a study that profiled luminal A breast cancer subtype (lumA BRCA) and matched controls, a disease in which metabolic pathways are known to be disrupted as a natural consequence of disease development. From the Cancer Genome Atlas (TCGA) we identified 60 individuals for whom expression profiling had been performed on both luminal A tumor and matched normal adjacent tissue (Cancer Genome Atlas Research Network et al., 2013). We downloaded the corresponding uniformly normalized and standardized gene expression data from Recount3 (Wilks et al., 2021) and inferred a pair of tumor and normal PANDA gene regulatory networks (see Methods). PANDA is a regulatory network inference method that integrates transcription factor motif and protein-protein interactions prior knowledge with gene expression to create bipartite transcription factor-to-gene regulatory networks (Glass et al., 2013). We transformed the bipartite graphs to unipartite by projecting them to a pair of gene-by-gene matrices of size 25410 *×* 25410 that represent gene co-regulation networks. We then performed graph differential analysis on these networks with n2v2r and DeDi, selecting the top 2% (roughly 500) genes from each method to perform over-representation analysis using the GOBP (Gene Ontology, Biological Process) database (Ashburner et al., 2000). As described previously, we ran n2v2r with Borda aggregation (n2v2r b) for dimensionalities *d* = {4, 6, 8, 10, 12, 14, 16, 18, 20, 22, 24} with Euclidean and cosine distances (which is our default setting in all analyses). The runtime for n2v2r was approximately 5 minutes on a c5.12xlarge AWS EC2 instance. Figure 3 shows the top 30 FDR-adjusted GO processes from n2v2r, with bold font indicating those found also by DeDi (Supplementary Material Figure S7).

**Figure 3.**
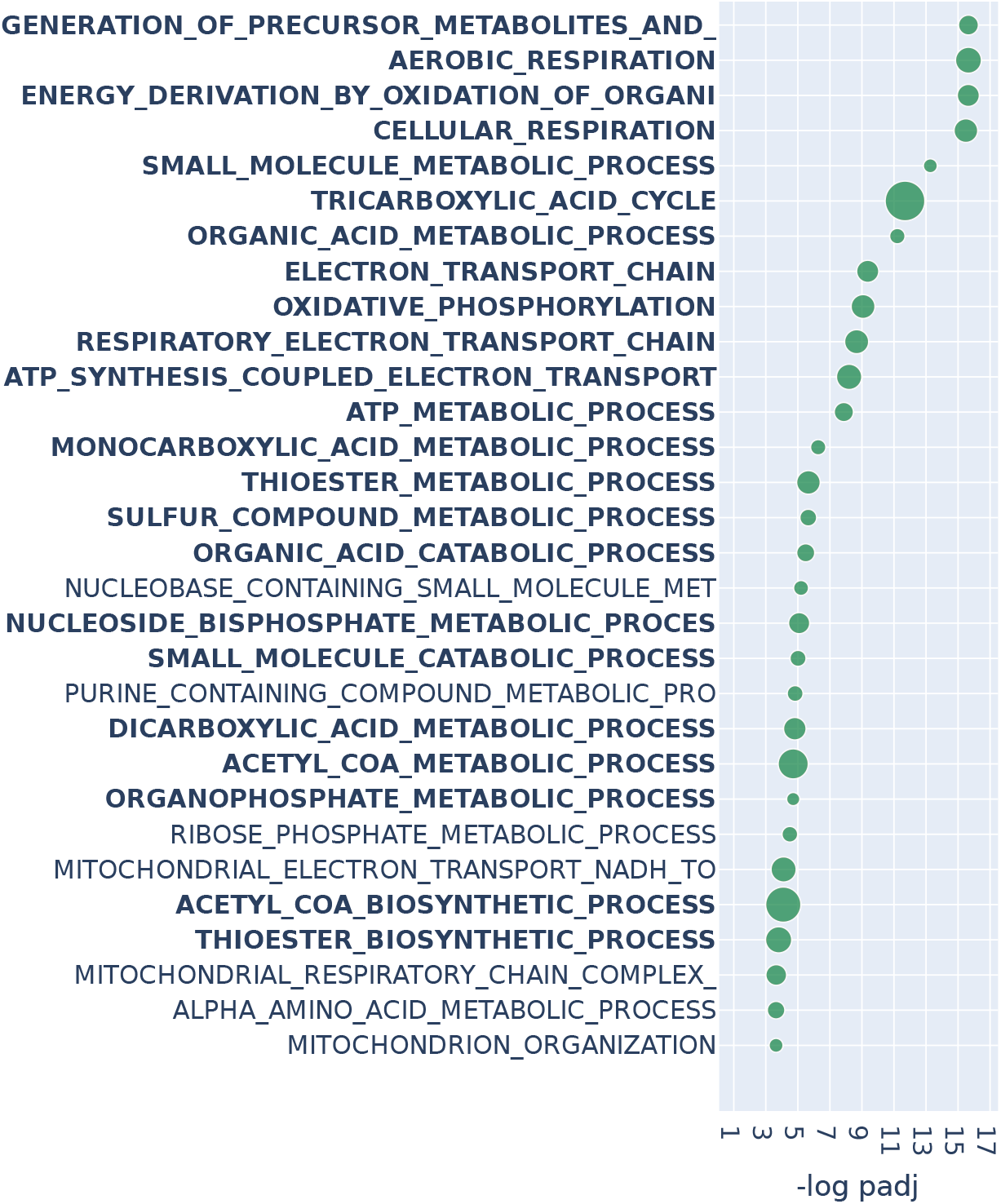
Over-representation analysis with GOBP (Gene Ontology, Biological Process) annotations on BRCA TCGA. We consider the top 2% of genes (roughly 500) of n2v2r on BRCA TCGA data when comparing luminal A tumor samples against matched adjacent normal samples using PANDA regulatory networks. We present the top 30 results ranked by adjusted *p*-values that pass a 0.1 FDR cutoff (equivalent to − log padj = 1). Bold indicates GOBP terms common with DeDi when performing the same analysis. The size of the points indicates the overlap of the input gene set with the annotated GO process. Long GOBP term names have been truncated.

Many of the GOBP terms we found to be significant are associated with metabolic processes, concordant with the hypothesis that a driver of luminal A breast cancer is the alteration of metabolism (Prat et al., 2015; Lunetti et al., 2019). Specifically, we found processes associated with the mitochondria, including oxidative phosphorylation (OXPHOS), consistent with reports that luminal A breast tumors exhibit increased OXPHOS activity relative to basal-like breast cancer and other subtypes (Cappelletti et al., 2017; Ortega-Lozano et al., 2022)—and that OXPHOS activity is predictive of outcomes in estrogen receptor (ER) positive breast cancer (El-Botty et al., 2023; Koc et al., 2022). Changes in OXPHOS and mitochondrial functions more generally can alter how breast cancer cells adapt their metabolism to support rapid growth, survive under various conditions, and evade therapeutic interventions (Avagliano et al., 2019).

Increased regulatory connectivity of metabolically relevant genes for lumA can also be seen in the network comparison in Supplementary Material Figures S8, S9. For example, the NDUFA9 and SDHA genes are common between multiple significant GOBP terms. NDUFA9 is a component of complex I in the respiratory chain, and its downregulation has been reported to promote metastasis by mediating mitochondrial function (Stroud et al., 2012; Li et al., 2015). SDHA, which is known to be a tumor suppressor gene, encodes one of the four subunits of the succinate dehydrogenase (SDH) enzyme, which plays a critical role in both OXPHOS and the Krebs cycle (Rutter et al., 2010); SDHA has also been identified as a marker for breast tumor progression and prognosis (Kim et al., 2013).

In contrast, differential gene expression analysis using DESeq2 followed by over-representation analysis for GOBP terms found only enrichment for general development processes that do not speak to the mechanisms that are altered in a disease-relevant manner (Supplementary Material Figure S10). The far deeper and more biologically relevant identification of specific GOBP terms demonstrates the importance of network analyses compared to simple differential expression, as we have reported previously in other contexts (Weighill et al., 2021).

#### 3.3.2 Monitoring Cell-cycle Transitions in Single-cell Data

We investigated cell-cycle transitions in single HeLa cells using n2v2r and DeDi. Cells undergo growth transition between four states: G1→S→G2→M when they grow and divide as part of the cell cycle. G1 is the first growth phase, followed by S, the synthesis phase, during which cells replicate their DNA. Cells then enter a second growth phase G2, which is followed by mitosis (M) in which cells divide into two daughter cells, after which the process repeats. We used combined data from wild-type and AGO2 knock-out HeLa S3 human cell lines comprising 1477 unsynchronized cells (Schwabe et al., 2020). We integrated the data using Seurat (Hao et al., 2021) and aggregated single-cells to create “metacells” that represent each of the four cell-cycle states. For these, we constructed four topological overlap networks of dimension 2420 *×* 2420 using hdWGCNA (Morabito et al., 2023) (see Methods). We applied n2v2r jointly on all networks using the sequential strategy of computing pairwise rankings of networks against the next in the cycle. To facilitate validation, we studied the well-known G1→S and G2→M transitions for which the Reactome database contains explicitly annotated pathways (Gillespie et al., 2021). Runtime on a c5.12xlarge AWS EC2 instance was roughly 10 seconds.

An overview of the data integration and network inference process is given in Figure 4a. Figure 4b, shows the top 30 FDR-adjusted gene set enrichment analysis (GSEA) results based on Kolmogorov–Smirnov testing for over-representation of Reactome pathways (see Methods). The green boxes indicate pathways that explicitly refer to the G1→S transition, and bold text indicates results also found using DeDi (corresponding enrichment results can be found in Supplementary Material Figure S11). We see that n2v2r identifies at least three exclusive pathways that are related to G1→S while DeDi finds none. We also observe that both methods identify common pathways associated with the G2→M transition. Panel c shows the top 30 GSEA results for the G2→M transition, with purple boxes indicating Reactome pathways explicitly annotated to this transition. We see that n2v2r finds pathways related to G2→M and with large enrichment scores, while DeDi finds no significant results.

**Figure 4.**
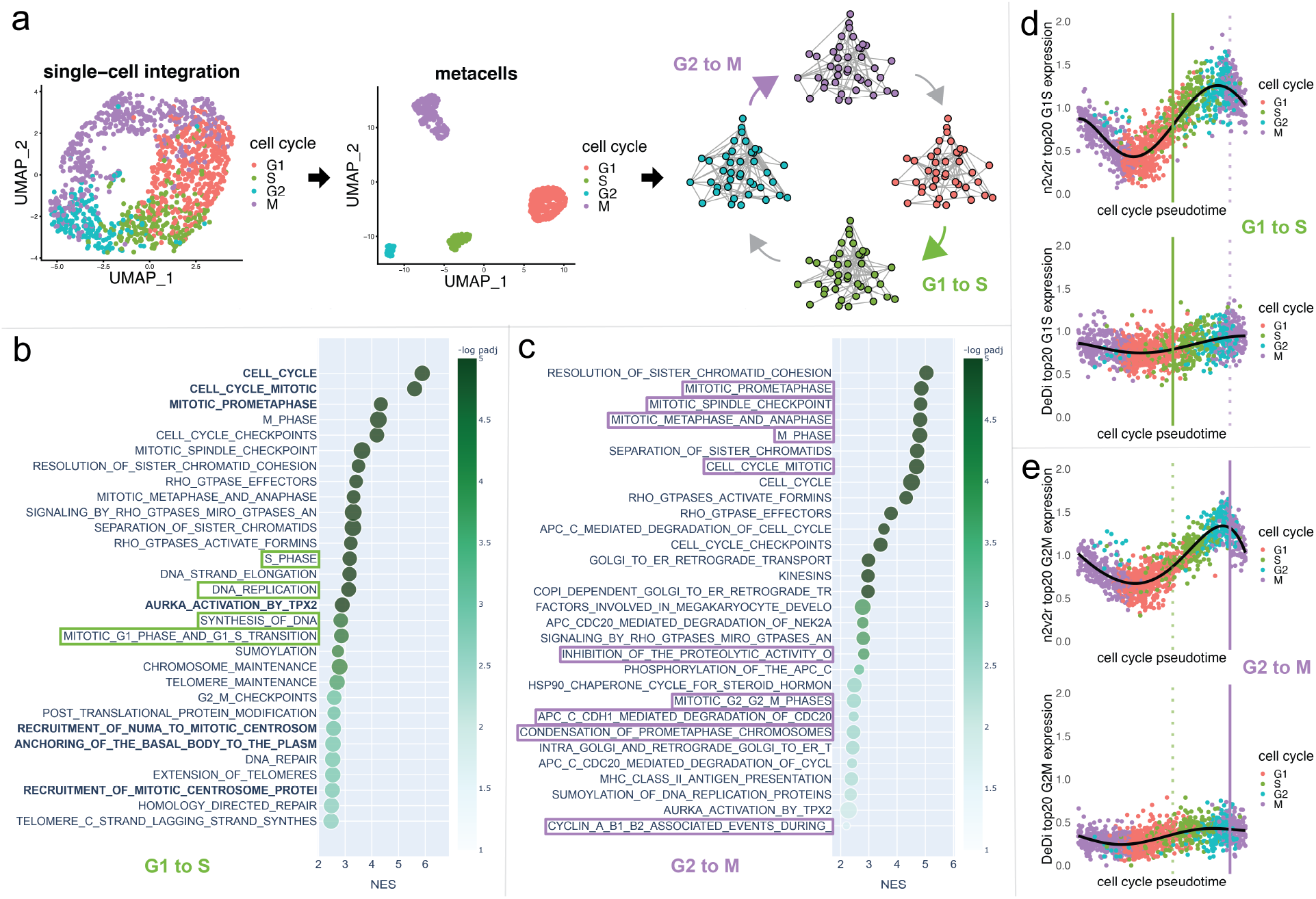
HeLa single-cell cell-cycle transitions analysis. (a) The HeLa S3 cell line was processed using Seurat and cells were aggregated to create metacells from which four co-expression networks corresponding to the cell-cycle phases G1, S, G2, and M were generated using hdWGCNA; subsequent analyses were limited to the G1 to S (green) and G2 to M (purple) transitions. (b) The top 30 results ranked by adjusted *p*-values (FDR 0.1 cutoff, equivalent to − log padj = 1) from gene set enrichment analysis (GSEA) based on Reactome annotation for genes found after applying n2v2r to the G1 to S transition are shown. The x-axis represents the normalized enrichment score (NES), and the size of the points indicates the overlap of the leading gene sets with the Reactome pathways. (c) GSEA results for the G2 to M transition; long pathway names have been truncated. (d) The average scaled and normalized expression for the top 20 genes from n2v2r (top) and DeDi (bottom) for the G1 to S transition. The x-axis represents the cell cycle pseudotime ordering of the cells using Revelio. The vertical lines represent the 0.9 quantile of pseudotime for G1 (left, green) and G2 (right, purple, dotted) cells, acting as proxies for the time the transitions occur. The black curve is a fifth-order polynomial fit to the data. (e) Similar pseudotime plots for the G2 to M transition.

Since the analysis here represents a time course captured in individual cells, we wanted to understand the overall profiles of genes found by n2v2r and DeDi during cell-cycle phase transitions. Figures 4d and e show the average gene expression of the top 20 genes in all cells identified by n2v2r (top) and DeDi (bottom) for the two transitions; cells are ordered according to cell-cycle pseudotime assigned using Revelio (Schwabe et al., 2020). It is interesting to note that the number of unique molecular identifiers (nUMI), corresponding to the number of distinct mRNA transcripts, increases with pseudotime and then drops after G2, which is where cell division occurs (Supplementary Material Figure S12). Given the processes active during the cell cycle, we expect the number of transcripts to track with marker genes that indicate the cell-cycle phase.

In panel d, with marker genes selected for the G1→S transition, we see that n2v2r identifies genes with a steep transition between these phases. In panel e, for the G2→M transition, n2v2r again retrieves genes that sharply decline in expression after entry into M, correctly capturing the point of mitosis. In contrast, the DeDi genes identified for both transitions were lowly expressed, lacking marker behavior, and thus failed to capture relevant cell-cycle processes.

#### 3.3.3 Sex Differences in Lung Adenocarcinoma

Lung cancer manifests differently in males and females, where both prognosis and response to therapy differ between the sexes, although mechanistic causes remain largely unknown (Legato et al., 2016; Ö zdemir et al., 2018; Reddy and Oliver, 2023; Silveyra et al., 2021). We applied n2v2r and DeDi to sex-specific gene coexpression networks from the lung adenocarcinoma (LUAD) TCGA cohort. We used the weighted gene co-expression network analysis (WGCNA) R package (Langfelder and Horvath, 2008) to construct two sex-specific 26000 *×* 26000 co-expression networks (see Methods). The runtime for n2v2r was approximately 1 minute on a c5.12xlarge AWS EC2 instance. We excluded Y chromosome genes from the rankings of both methods, as they often reflect inherent sex differences rather than disease-specific processes.

Figure 5 shows the top 30 enriched KEGG pathways based on gene set enrichment analysis using the ranked list from n2v2r; pathways also found using DeDi are shown in bold (Supplementary Material Figure S13). There was a 60% overlap (18 out of 30) between the enriched pathways identified using n2v2r and DeDi. Not surprisingly, many of the common pathways are related to immune processes, such as the T cell receptor and Toll-like receptor signaling, which are known to be sex-biased in LUAD (Klein and Flanagan, 2016). In addition, common pathways such as the spliceosome, endocytosis, ubiquitin-mediated proteolysis, and cell cycle, correspond to sex-biased cellular processes (Rubin et al., 2020; Rubin, 2022).

**Figure 5.**
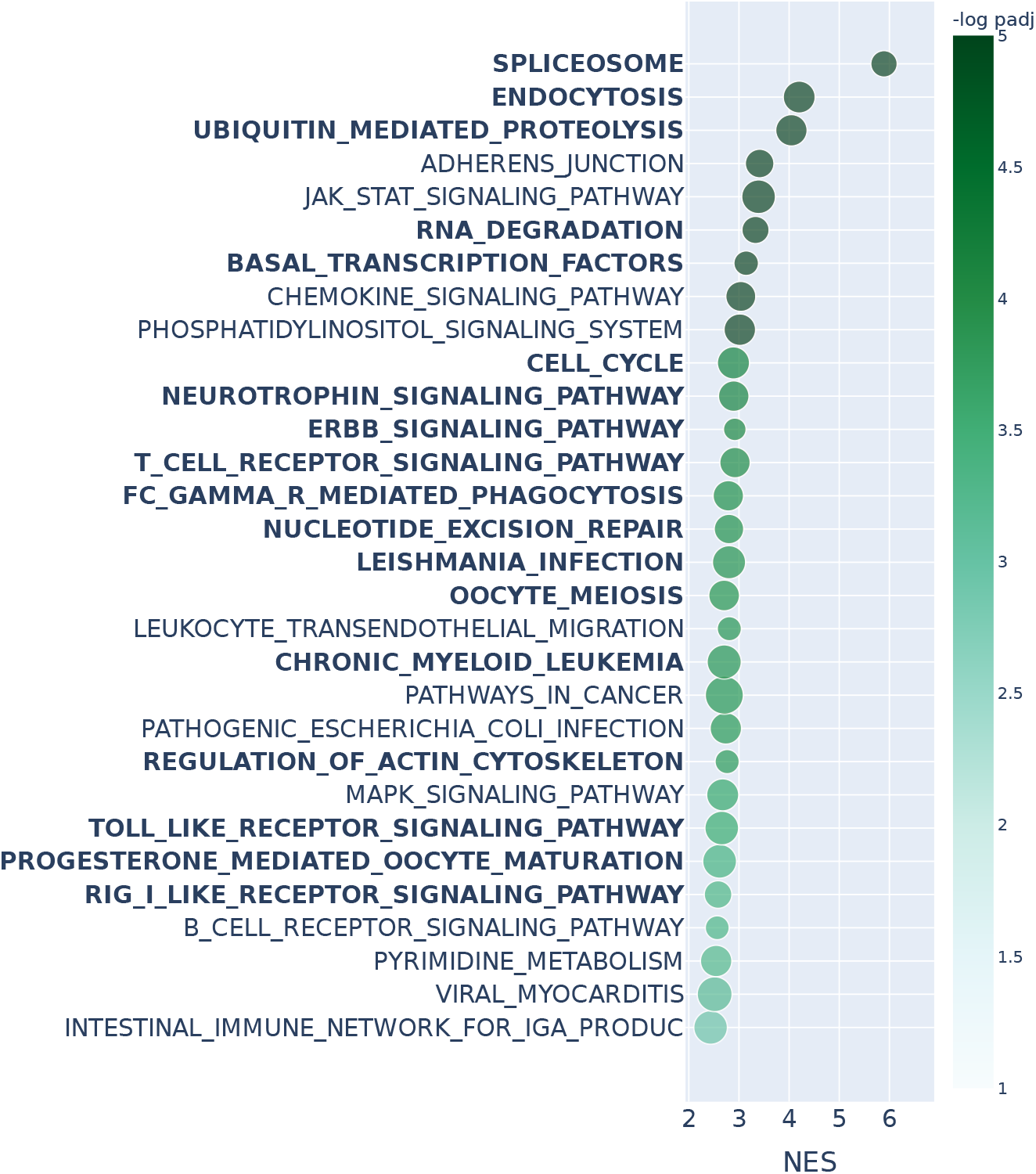
Gene set enrichment analysis with KEGG pathway annotations and Kolmogorov–Smirnov test on TCGA LUAD co-expression networks. We consider the entire n2v2r ranking of sex-specific differences found in WGCNA co-expression networks inferred for TCGA LUAD male and female samples. The top 30 results passing a 0.1 FDR cutoff (equivalent to − log padj = 1) and ranked by adjusted *p*-values are shown. Bold indicates pathways common with DeDi when performing the same analysis. The x-axis represents the normalized enrichment score, and the color represents the value − log padj. The size of the points indicates the overlap of the leading gene sets with the pathways. Long pathway names have been truncated.

The enriched pathways that are exclusive to the DeDi ranking include many related to neurodegenerative disorders, other cancer types, and broad cellular pathways such as ribosome and DNA replication (Figure S13)—pathways that provide little insight into sex-specific LUAD mechanisms. In contrast, the pathways found exclusively when using n2v2r include specific cancer-related processes such as the MAPK and phosphatidylinositol pathways that are known to be regulated by sex hormones and whose disruption has been shown to be linked to non-small cell lung cancer cell propagation and tumor growth (Fuentes et al., 2021). The n2v2r pathways also include B-cell receptor signaling, with B-cell related genes reported as significantly upregulated in the LUAD tumors of females, consistent with reports that LUAD tumors in females harbor a greater number of infiltrating B-cells (Klein and Flanagan, 2016; Li et al., 2023).

Among the pathways found only using n2v2r is the JAK/STAT signaling, a cellular communication pathway involved in the regulation of various physiological processes, including immune responses and cell growth and differentiation; the JAK/STAT pathway that has been identified as playing a role in lung cancer development and progression (Thomas et al., 2015; Hu et al., 2021). To understand the drivers of sex bias in this pathway, we identified the subset of genes from the n2v2r ranking that weighed most heavily in determining pathway to be significantly sex-biased. We plotted the subgraph of **A**^female^–**A**^male^, focusing on those leading genes as central nodes (with larger font) and the top 500 connections (Figure 6a). Node size reflects the degree difference, and color denotes whether the gene is significantly differentially expressed between males and females based on DESeq2 tabular analysis (Love et al., 2014). Because there are clinically documented differences between the sexes in chemotherapy response, we used the PRISM (Yu et al., 2016) and GDSC (Yang et al., 2012) databases, each of which contains results of small molecule drug treatments of LUAD cell lines derived from male and female donors. We found significant sex-based differences for compounds targeting genes in the JAK/STAT signaling pathway (see Methods). Figure 6b shows that inhibitors targeting five JAK/STAT pathway genes, EP300, JAK1, JAK2, TYK2, and AKT2, exhibit significant sex-biased responses. For JAK1, JAK2, TYK2, and AKT2, we observed a strong female bias in correlation network connectivity (only red edges), which corresponded with inhibitory drugs being more effective for females; inhibitors of EP300 produced a better response in males.

**Figure 6.**
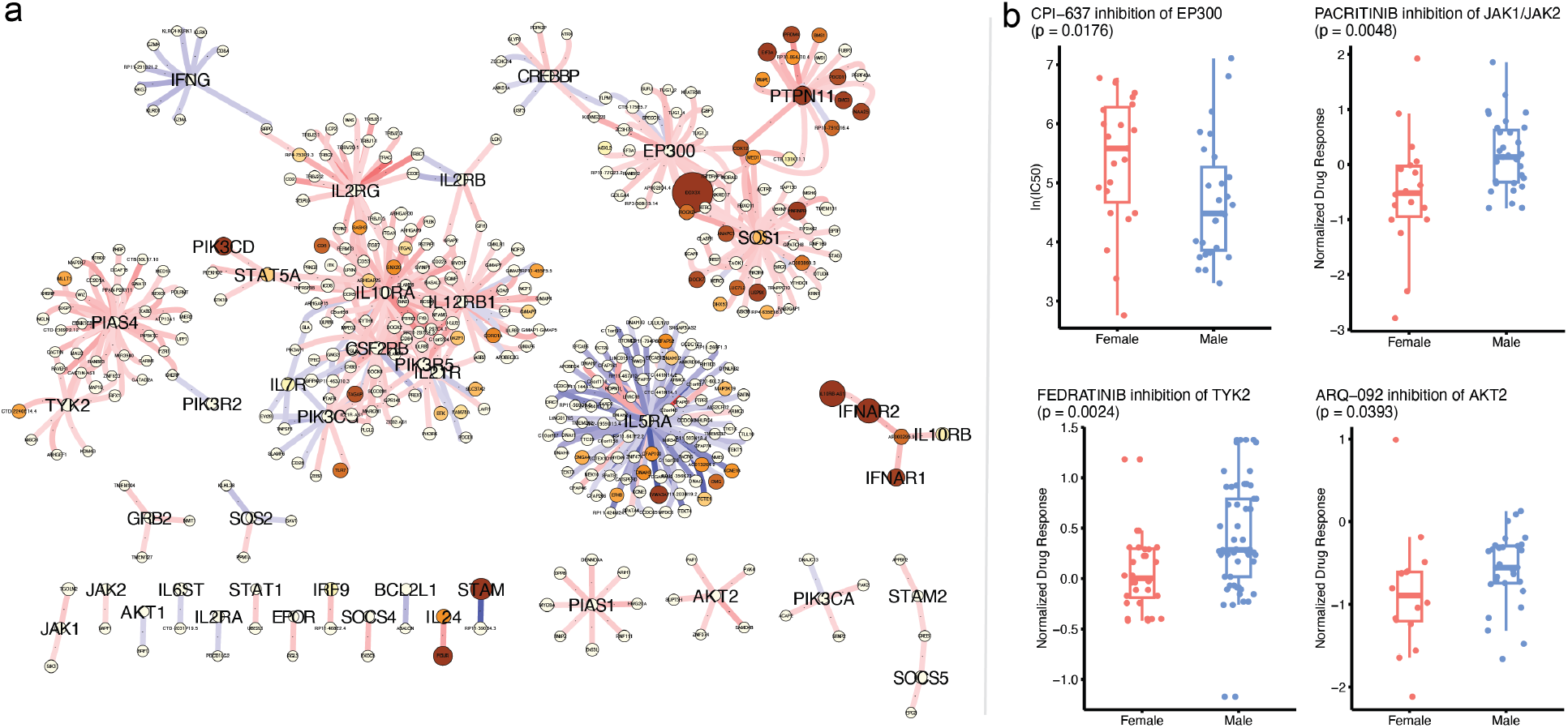
Probing the leading genes of the JAK/STAT signaling pathway for sex differences. (a) The top 500 edges of the differential network **A**^female^–**A**^male^ centered on the n2v2r leading genes of the JAK/STAT pathway as returned by GSEApy, shown here with larger fonts. Red edges indicate greater co-expression in females, and blue edges indicate greater co-expression in males. The size of the nodes is the absolute degree difference. Nodes are colored according to the adjusted *p*-value from differential gene expression analysis with DESeq2: white is not significant, yellow to red is significant between 0.1 and 0. (b) Significant sex-biased drug responses (lower more sensitive) by inhibiting JAK/STAT genes in LUAD cell lines that are not differential in node degree and gene expression. The EP300 is from the GDSC repository, the rest from PRISM.

One additional gene, IL5RA, also exhibited substantial sex-biased differences in correlation patterns (Figure 6). The IL5RA protein is involved in the IL-5 signaling pathway, which plays a key role in regulating immune responses and the activation of eosinophils. IL5RA has been implicated in multiple pulmonary diseases, including COPD and asthma, as well as LUAD (Fan et al., 2021; Sibille et al., 2022). Little is known, however, regarding the role of IL5RA in LUAD sex differences. A recent meta-analysis of GWAS data found IL5RA to be significantly sex-biased in lung epithelial tissue (Gautam et al., 2019), consistent with our observation of a male bias in the differential network (more blue edges). This suggests that sex-specific differences in the immune microenvironment, including the presence of specific immune cells such as eosinophils, may influence the course of the disease and possibly influence response to immunotherapy checkpoint inhibitors in a sex-biased manner.

Lastly, the results for differential gene expression analysis with DESeq2 (Supplementary Material Figure S14) primarily identified differences in well-known, sex-biased metabolic processes, finding few relevant cancer-related processes (Krumsiek et al., 2015).

Overall, the results from the three biological applications we considered show that node2vec2rank can effectively compare high-dimensional networks to produce biologically meaningful results and clearly demonstrate its advantages over alternative methods such as DeDi and tabular analysis.

## 4 Discussion

Differential expression analysis tools such as DESeq2 have demonstrated their value in identifying genes whose expression levels change between conditions. However, examining individual genes alone, or as collections of individual objects, overlooks the complex, higher-order processes that define phenotypes and govern transitions between biological states. Network-based methods overcome this limitation by modeling genome-wide relationships—correlation networks built with WGCNA or regulatory networks inferred using PANDA, for instance, can capture intricate dependencies among genes. Yet, comparing such networks across phenotypes has been an open problem: network graphs vary greatly in size and structure, and their higher-order associations are difficult to align. Methods like Degree Difference ranking (DeDi) can identify nodes whose connectivity shifts between conditions, but these emphasize properties of individual nodes and so fail to fully exploit the rich, multiscale organization embedded in biological networks.

We developed node2vec2rank (n2v2r), a multi-layer node embedding framework that takes advantage of the rich representational power of graphs to enable rigorous network differential analysis and overcome the limitations of feature-based methods. Unlike bag-of-features methods that rely on user-defined summary statistics, n2v2r learns node representations directly from the data, capturing latent structures that extend beyond simple connectivity measures such as node degree. Built on the unfolded adjacency spectral embedding (UASE) algorithm and singular value decomposition, n2v2r offers a statistically sound, efficient, stable, and interpretable way of ranking differential nodes across multi-layer weighted networks. The resulting lists of ranked nodes can be seamlessly integrated into established downstream analyses, including techniques such as pre-ranked gene set enrichment analysis.

Applying n2v2r to data from three diverse biological systems, we showed that it could identify key sets of genes representing relevant biological functions—functions that were not detected through either differential expression analysis with DESeq2 or graph-based degree difference analysis with DeDi. For luminal A breast cancer, we used bulk gene expression data from the Cancer Genome Atlas project (TCGA) with PANDA to derive co-regulatory networks. Using these, we showed that both n2v2r and DeDi rankings were able to identify key metabolic processes such as oxidative phosphorylation (OXPHOS), surpassing DESeq2 applied to the expression data.

We performed a cell-cycle analysis with gene co-expression networks derived from single-cell expression data. There, we found that n2v2r outperformed DeDi by identifying multiple biological processes known to be activated during the respective cell-cycle phases and found marker genes that exhibit meaningful oscillatory expression patterns over the entirety of the cell cycle.

We also investigated sex differences in co-expression networks derived from TCGA lung adenocarcinoma (LUAD) data. Using n2v2r with these networks, we found pathways known to be sex-biased in LUAD, including B-cell-associated processes and other immune responses relevant to understanding LUAD development, progression, and response to cancer immunotherapy. We dissected the JAK/STAT signaling pathway, which was only identified when using n2v2r, and we found genes that were neither differentially expressed nor exhibited differential node degree, but had plausible connections to cancer. This is significant as drugs that inhibit the corresponding proteins in lung cancer cell lines exhibit sex-biased responses, meaning that n2v2r recovered actionable regulatory patterns missed by standard analysis methods. We also found evidence for a sex-biased role of IL5RA in LUAD, a result supported by its involvement in several pulmonary diseases and the significant sex bias in lung epithelial tissue that had been reported in a genome-wide association meta-analysis.

One challenge in developing n2v2r was the choice of embedding dimensionality. It is common for machine learning methods to use a large number of dimensions (such as 64, 128, 256, or more), but measuring distances in these embedding spaces tends to become meaningless as the dimensions grow. The exact number of dimensions in which this occurs, especially for graphs with different characteristics, is difficult to determine. The Borda aggregation helps to average and stabilize performance, which is beneficial in cases of misspecification. As a default, we opted for a moderate number of dimensions, ranging between 4 to 24 with an increment of 2, and for each dimension, we used cosine distance and Euclidean distance, giving a total of 22 distance-based rankings. This choice of embedding dimensions yielded consistently good results in each of the three applications of n2v2r in biological data. We recognize that our choice of embedding dimensions is somewhat arbitrary; a data-driven alternative would be to consider the singular values and the amount of variance explained, and to use a window centered on the dimension corresponding to the elbow point in the scree plot.

Overall, node2vec2rank (n2v2r) provides a theoretically grounded and practically effective framework for comparing networks, and its performance has been validated across diverse biological applications. While our analysis focused on unipartite networks, we anticipate that the method could be extended to multipartite networks, which more accurately capture processes such as gene regulation. Spectral embedding is already applicable to single multipartite networks (Modell et al., 2022), but extending it to multiple networks remains an open challenge. Determining which network is most affected by differences in nodes or pathways could help infer directionality between related networks; for instance, one could interpret a larger difference in node embedding norms as indicating a more active node in the comparison. Further extensions incorporating nonlinear interactions, local neighborhood effects, and node and edge features could reveal more complex differential patterns, fully leveraging the richness of network structure.

Although we assessed n2v2r in analyses of biological systems, there is nothing in its derivation or properties that would prevent its use in other graph comparison tasks. Given its robustness and stability, as well as its adherence to the veridical data science principles of predictability, computability, and stability (PCS), we can see potential for broad applications of n2v2r in other disciplines.

## 5 Methods

### 5.1 Fitting Node2vec2rank with UASE

#### 5.1.1 Multi-Layer Latent Position Model

We assume there exists a latent space *Ƶ* in which the latent position 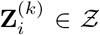 dictates how node *i* forms edges in graph **A**^(*k*)^. The weighted multi-layer latent position model assumes that the weight of an edge (*i, j*) in **A**^(*k*)^ depends only on the latent positions of nodes *i, j*. For all *k* ∈ [*K*] and *i < j*, we have,

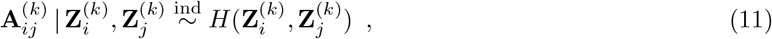

where *H* is a symmetric real-valued distribution function, *H*(**Z**_1_, **Z**_2_) = *H*(**Z**_2_, **Z**_1_) for all **Z**_1_, **Z**_2_ ∈ *Ƶ*.

Under an approximate low-rank adjacency matrix assumption (see Appendix), valid for many real-world networks (Udell and Townsend, 2019), there exists a map *ϕ*_*k*_ : *Z* → ℝ^*d*^ representing the latent positions in a *d*-dimensional space such that,

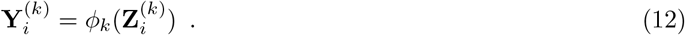

Under this assumption, this general model encompasses a range of graph types extending the weighted generalized random dot product graph (Gallagher et al., 2023). For unweighted networks, *H* is a Bernoulli distribution with some link probability function 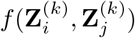, and the latent position model is a special case of the multi-layer random dot product graph (Jones and Rubin-Delanchy, 2020; Gallagher et al., 2021). This includes the stochastic block model as well as mixed membership and degree-corrected variants.

#### 5.1.2 Unfolded Adjacency Spectral Embedding

Given weighted adjacency matrices **A**^(1)^, …, **A**^(*K*)^ ∈ ℝ^*n×n*^, we construct the unfolded adjacency matrix using column concatenation,

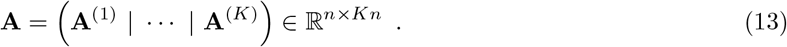

Given embedding dimension *d*, we compute the *d*-truncated singular value decomposition (SVD) of 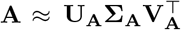, where **U**_**A**_ ∈ ℝ^*n×d*^, **V**_**A**_ ∈ ℝ^*Kn×d*^ are truncated left and right singular vectors, and **Σ**_**A**_ ∈ ℝ^*d×d*^ is the diagonal matrix with the *d* largest singular values. Using row concatenation to the right singular vectors corresponding to the *K* graphs, we have,

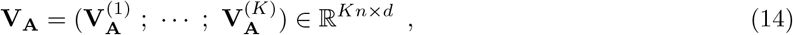

and the UASE for graph **A**^(*k*)^ is defined by,

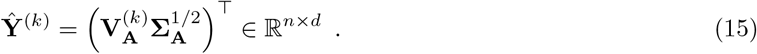

#### 5.1.3 Assumptions and Proofs

In this section, we outline the necessary model assumptions and provide a sketch proof of Theorem 1, covering the main steps rather than going deeply into technical details. The proof follows a similar argument to the consistency proof from the original generalized random dot product graph model publication by Rubin and colleagues (Rubin-Delanchy et al., 2022). This proof was separately modified to accommodate weighted networks (Gallagher et al., 2023) and multi-layer networks (Jones and Rubin-Delanchy, 2020). Combining these two sets of adjustments demonstrates the consistency of unfolded adjacency spectral embedding.

We introduce three assumptions on the weighted multi-layer latent position model to enable sensible node embeddings into a low-dimensional space.

##### Assumption 1

(Low-rank expectation). *There exist maps ϕ* : *Ƶ*^*K*^ → ℝ^*d*^ *and ϕ*_*k*_ : *Ƶ* → ℝ^*d*^ *for k* ∈ [*K*] *such that*,

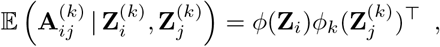

*Where* 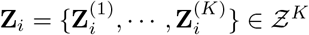.

Assumption 1 is not a practical restriction on the multi-layer graph model, as many real-world networks have low rank (Udell and Townsend, 2019). We define t he o utput o f t he l atent p osition m aps from Assumption 1 by the matrices **X** and **Y**^(*k*)^ with rows given by,

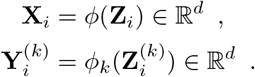

One possible choice for the maps *ϕ*_*k*_, which defines a choice for the map *ϕ*, is based on the mean unfolded adjacency matrix **P** = E(**A**). The low-rank assumption states that the *d*-truncated SVD 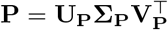 is exact. Using these left and right singular vectors, we can define a canonical choice for the maps,

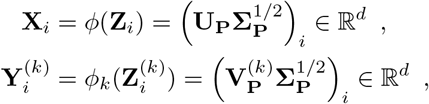

where 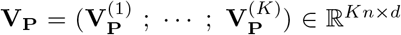 divides the right singular vector **V**_**P**_ into blocks corresponding to the *K* graphs using row concatenation. Other choices for the maps *ϕ*_*k*_ are possible, but these ultimately represent orthogonal transformations of the embeddings **Y**^(*k*)^. This will get absorbed into the orthogonal transformation **W** ∈ 𝕆 (*d*) in Theorem 1.

##### Assumption 2

(Sub-exponential tails). *There exists a constant α >* 0 *such that*

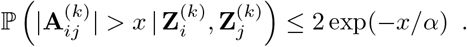

Assumption 2 states that the edge weights are bounded with high probability. This is necessary to prevent very large edge weights in the networks, which would otherwise ruin the structure in an embedding. It is satisfied trivially for bounded distributions, such as the Bernoulli and beta distributions, as well as many other standard distributions, including the exponential distribution and the Gaussian distribution.

##### Assumption 3

(Singular values of **P**). *The d non-zero singular values of the mean unfolded adjacency matrix* **P** = 𝔼 (**A**) *satisfy*

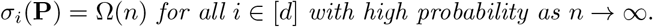

Assumption 3 is usually written as a condition on the distribution of the latent positions 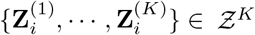 controlling how the matrices **X**^⊤^**X** and **Y**^(*k*)⊤^**Y**^(*k*)^ deviate from their corresponding second moment matrices (Rubin-Delanchy et al., 2022; Jones and Rubin-Delanchy, 2020; Gallagher et al., 2023). However, the goal of this assumption is always the same: regulating the growth of the singular values of the matrix **P** as *n* → ∞. In our analysis, we directly assume this condition, rather than introducing distributions for the latent positions 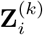.

##### Proof of Theorem 1.

We now outline a sketch proof of Theorem 1 under model Assumptions 1, 2, and 3.

*Proof outline*. The starting point for proving the theorem involves considering the difference between the unfolded adjacency matrix **A** and its unfolded mean matrix **P**. Under Assumption 2, the spectral norm of their difference has the following asymptotic bound,

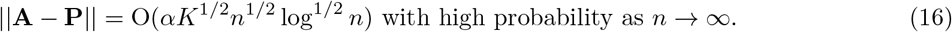

This is a consequence of the sub-exponential version of the matrix Bernstein inequality (Theorem 6.2, Tropp (2012)). From this property, both the singular value matrices **Σ**_**A**_ and **Σ**_**P**_, and the subspaces spanned by **V**_**A**_ and **V**_**P**_, are approximately equal. The first is a consequence of Weyl’s inequality,

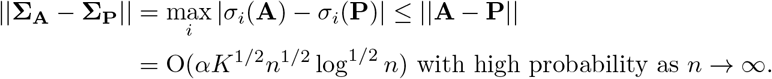

The second is derived from the Davis-Kahan sin *θ* theorem (Yu et al., 2015) using Assumption 3 about the size of *σ*_*d*_(**P**),

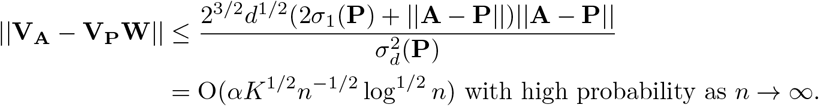

The orthogonal matrix **W** here solves the one-mode orthogonal Procrustes problem that aligns the left and right singular vectors of the matrices **A** and **P**,

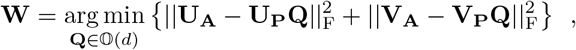

where || · ||_F_ denotes the Frobenius norm. Using this orthogonal matrix **W**, we can compare how the UASE over all graphs 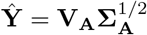 behaves in relation to the canonical choice 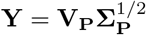 defined by the maps *ϕ*_*k*_. The proof uses the decomposition,

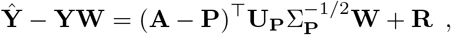

for some residual matrix **R** ∈ ℝ^*n×d*^, which implies that, for all *k* ∈ [*K*],

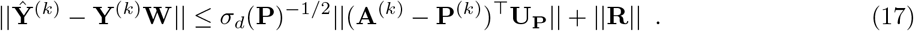

Using the asymptotic behavior of ||**A** − **P**|| from Equation 16, the first term is controlled asymptotically by the bound,

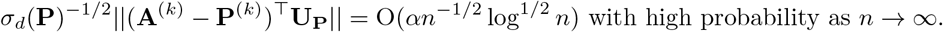

The majority of the proof is to show that the residual matrix **R** is well-behaved, starting from the spectral norm properties described above. This is a technical exercise in matrix perturbation using the existing techniques from the proofs for binary, weighted, and multi-layer graphs (Rubin-Delanchy et al., 2022; Gallagher et al., 2023; Jones and Rubin-Delanchy, 2020). Applying the same approach in this setting gives

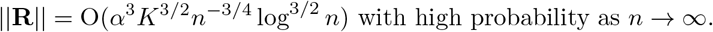

This is the dominating term in Equation 17, meaning that

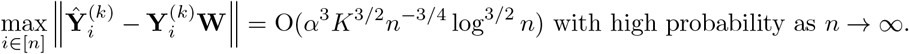

This controls the rate of convergence, although the statement in Theorem 1 is only concerned that this converges to zero as *n* → ∞.

##### Proof of Theorem 2

*Proof*. The proof uses invariance under orthogonal transformation and the triangle inequality of Euclidean and cosine distances,

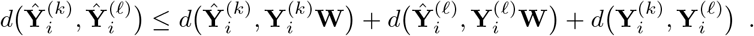

Alternatively, we could have started considering the distance between the latent positions and derived a similar inequality,

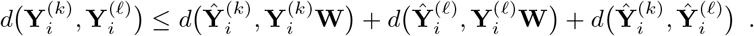

Together, these give the following inequality for the maximum possible error over all nodes for any distance functions invariant under orthogonal transformation,

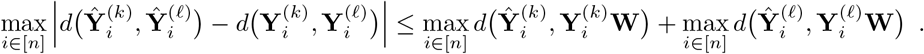

For Euclidean distance, Theorem 1 states that the two terms on the right-hand side tend to zero with high probability as *n* → ∞ giving the first result. For cosine distance, if a sequence of points converges with respect to Euclidean distance to a non-zero limit, then the sequence also converges with respect to cosine distance. Therefore, if both 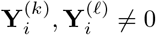, Theorem 1 states that the two terms on the right-hand side tend to zero with high probability as *n* → ∞ as given by the second result.

### 5.2 Simulated Network Generation

We generated pairs of binary graphs **A**^(1)^, **A**^(2)^ ∈ {0, 1}^*n×n*^ following two stochastic block models with *m* communities and community probability matrices **B**^(1)^, **B**^(2)^ ∈ ℝ^*m×m*^. Each node is assigned to one of the *m* communities uniformly at random, the same in both graphs 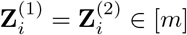. For convenience, we drop the unnecessary superscript for community assignments.

First, we generated the adjacency matrix **A**^(1)^ by independently sampling Bernoulli edges for each pair of nodes according to their community assignments,

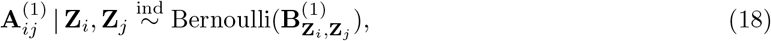

as given by the multi-layer latent position model in Equation 11. However, the method for generating the adjacency matrix **A**^(2)^ will differ from this latent position model to provide a better test of the n2v2r framework.

To generate **A**^(2)^, we adjusted the first graph to satisfy the correct community probabilities given by **B**^(2)^,

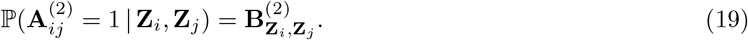

We considered two possible cases for the pair of nodes *i* and *j*:

- If 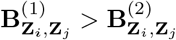, we deleted the edge (*i, j*) if it exists with probability,

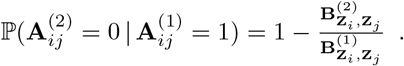
- If 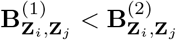, we add the edge (*i, j*) if it does not exist with probability,

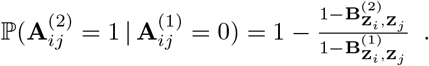

In all other cases, the adjacency matrices remain the same. Under this sampling method, Equation 19 is satisfied.

### 5.3 Simulated Network Differences and Ground Truth Assignment

Both the degree-naive and random differences regimes were simulated by perturbing a randomly sampled community probability matrix **B**^(1)^ to create a second **B**^(2)^. For the degree-naive differences, we sampled a **B**^(1)^ uniformly at random with values ranging in [0, 0.5]. Then, to create **B**^(2)^, we “flipped” the first row vector and column vectors of **B**^(1)^ while maintaining 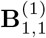 as is. The effect of this transformation is that the community represented by the first row and column will have its connections with the other communities reassigned, but the overall degree remains the same. An example of this operation with four communities is the following, highlighting in bold the elements to be flipped:

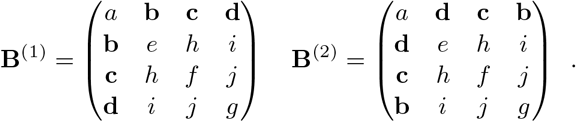

Finally, we added noise to the remaining elements of **B**^(2)^ with 50% probability following a normal distribution 𝒩 (0, ^0.5^*/*_10_).

For the second regime of random differences, we followed a similar scheme. First, we sampled a **B**^(1)^ uniformly at random with values ranging in [0, 0.5]. We then created **B**^(2)^ by completely re-sampling the first row and column of **B**^(1)^ with another set of values in [0, 0.5]. Then, we proceed to add noise to the remaining elements of **B**^(2)^ with 50% probability following a normal distribution 𝒩 (0, ^0.5^*/*_10_).

Note that the Gaussian noise can cause the altered community to fall in ranks, which would invalidate its ground truth assignment as the most changing. To overcome this, we introduce a rejection sampling scheme where we reject a pair of **B**^(1)^, **B**^(2)^ matrices if the community-wise Borda aggregated Euclidean, cosine, and Chebyshev distances between **B**^(1)^ and **B**^(2)^ do not rank the altered community as the one changing the most.

### 5.4 Single-cell Cell-Cycle Network Generation and Analysis

We used the single-cell data of Schwabe et al. (2020), GEO accession GSE142277, corresponding to 1477 cells from HeLa S3 wild-type and AGO2 knock-out (AGO2KO) cell lines (Schwabe et al., 2020). We used the Seurat v4.3.0.1 pipeline to remove cells with fewer than 200 genes detected or a mitochondrial ratio greater than 5%, leaving 1282 cells. We used *SCTransform* to regress out the two batches, wild-type and AGO2KO, leaving behind expression profiles that minimize the effect of the gene knock-out.

We then used hdWGCNA (Morabito et al., 2023) v0.2.26 to create gene correlation networks for metacells. We first created metacells using the function *MetacellsByGroups*, which performs KNN on the PCA space. We set the number of neighbors *k* = 20, set the maximum number of shared cells between two metacells to 10, and grouped the results into the four cell-cycle stages. We used this setting to obtain a reasonable number of metacells; 792 metacells (out of 1282) met these criteria. Using the functions *SetDatExpr* and *PlotSoftPowers* with default settings, we tested for soft-thresholds for each network, after which we chose 5 for all networks to avoid any imbalances; soft-thresholding affects the mean connectivity of the networks. We constructed four topological overlap matrices (TOM) using the function *ConstructNetwork*. The TOM is a similarity matrix based on the number of genes two genes have in common after applying a threshold. We dropped all genes that were not common to all networks, resulting in 2420 shared genes. For the plots in Figures 4d and e, we used the functions *getPCAData* and *getOptimalRotation* from Revelio v0.1.0 to retrieve the cycle pseudotime and order the cells.

### 5.5 Bulk Data Network Generation

Gene expression data for breast cancer (BRCA) and lung adenocarcinoma (LUAD) were collected by The Cancer Genome Atlas (Cancer Genome Atlas Research Network et al., 2013) and subsequently reprocessed and made available by the Recount3 project (Wilks et al., 2021) downloaded (02 Nov 2022). We used routine gene expression pre-processing protocols with these data: counts were logTPM transformed, and we removed genes that had 1 TPM or less in more than 90% of the samples. We also remove samples that had less than 30% tumor purity using the consensus method from (Aran et al., 2015). That left 1201 samples and 25687 genes for BRCA, and 574 samples and 26000 genes for LUAD.

For the BRCA luminal A (lumA) and adjacent normal comparison, we filtered the processed data for lumA patients who had adjacent normal tissue samples, leaving 60 individuals with both. We removed four genes that had constant values across samples, leaving 25683 genes. We used the 60 tumor samples and the 60 adjacent normal samples to compute two PANDA networks (Glass et al., 2013) with netZooPy v0.9.13 (Ben Guebila et al., 2023).

PANDA requires as input both a transcription factor (TF)-to-gene motif binding prior and a TF-to-TF prior protein-protein interaction (PPI) network. Following (Lopes-Ramos et al., 2020), we generated the TF-to-gene prior network with putative motif targets using FIMO (Grant et al., 2011). We downloaded the Homo sapiens TF motifs with direct or inferred evidence from the Catalog of Inferred Sequence Binding Preferences (CIS-BP) Build 2.0 (http://cisbp.ccbr.utoronto.ca). FIMO maps the position weight matrices (PWM) to the human genome (hg38), and we used only significant matches (threshold: *p* ≤ 10^−5^) occurring within the promoter regions of Ensembl genes (Gencode v39 (Frankish et al., 2020), downloaded from http://genome.ucsc.edu/cgi-bin/hgTables). The promoter regions were defined as [−750; +250] base pairs upstream and downstream of the transcription start site (TSS), respectively. This produced a prior network consisting of 997 transcription factors that targeted 61485 genes. For the TF PPI networks, we downloaded PPI data from the StringDB database (version 11.5) (Szklarczyk et al., 2021) using the associated Bioconductor package. We kept only the TFs present in the motif prior, and we normalized these by dividing each score by 1000, thus restricting the values between 0 and 1. We also transformed the PPI networks into symmetric networks with self-loops.

After running PANDA on both lumA cancer and control populations, we obtained two bipartite networks of 997 TFs and 25410 genes. We converted each PANDA bipartite network *W* to a gene-by-gene unipartite graph that represents co-regulation with the transformation *W* ^*T*^ *W* that creates an edge between every gene that has evidence of regulation by a common TF, resulting in two 25410 *×* 25410 networks.

For the LUAD male versus female comparison, we used Recount3 expression data from 238 and 277 male and female tumor samples, respectively. Using the downloaded gene expression data, we computed two WGCNA unsigned networks (version 1.72.1) with soft-threshold 10 using the functions *pickSoftThreshold* and *adjacency* in hdWGCNA.

### 5.6 Over-representation and Gene Set Enrichment Analysis

For over-representation and gene set enrichment analyses, we used the GSEApy Python package (Fang et al., 2022). With function *enrichr*, we performed over-representation analysis with a hypergeometric test to evaluate whether an input set of genes is significantly associated with specific biological processes or pathways. We also used the *prerank* function to perform gene set enrichment analysis using a Kolmogorov–Smirnov test to assess whether genes at the extremes of a ranked list of genes are associated with functional or pathway annotations. To determine significance, we used the unweighted version with 1500 permutations. For biological annotations, we used three gene pathway annotation databases, GOBP (Ashburner et al., 2000), KEGG (Kanehisa and Goto, 2000), and Reactome (Gillespie et al., 2021), versions v7.5.1 downloaded from the Molecular Signatures Database (MSigDB) (http://www.broadinstitute.org/gsea/msigdb/collections.jsp). We filtered pathways with fewer than 5 genes and with more than 1500 genes. We used an FDR cutoff of 0.1 and ranked results based on their adjusted *p*-values.

### 5.7 Differential Graph Plotting and Gene Expression Analysis

To plot the differential graphs in the TCGA BRCA and LUAD analyses, we used Cytoscape (Shannon et al., 2003). For differential expression analysis, we used the R package DESeq2 (Love et al., 2014) v1.36.0 with the original RNA-seq data (un-normalized counts) for the same genes and samples used in our network construction. Genes were ranked for analysis based on DESeq2 adjusted *p*-values.

### 5.8 LUAD Drug Responses

To validate predictions of n2v2r, we used drug response data from GDSC (Yang et al., 2012) and PRISM (Yu et al., 2016) for LUAD cell lines representing samples from both males and females. We compared drug response outcomes between the sexes, focusing on the most significant genes found in the JAK/STAT using n2v2r, but which were not differentially expressed. We performed a non-exhaustive search through these genes using the Wilcoxon rank sum test. For many genes, including IL10RA, IL2RA, PIAS1, PIAS4, and SOS2, there were entries in only GDSC or PRISM but not both. EP300 was significantly biased in GDSC but not in PRISM, while AKT2, JAK1, and JAK2 were significantly biased in PRISM but not in GDSC. TYK2 had no drug response in GDSC but was significantly biased in PRISM. IL5RA had no entries in GDSC and was not significantly biased in PRISM.

## 6 Reproducibilty Statement and Data Availability

Tool versions are provided when cited. Notebooks to recreate the bubbleplots and subgraphs in the analysis of the three biological datasets are available at https://github.com/pmandros/n2v2r; the YAML file reproduces the exact Python environment we used. The scripts we used to generate the single-cell, bulk, and simulated networks, as well as the drug responses, are available at Zenodo 10.5281/zenodo.10558426. Additionally, the same repository contains the biological networks and gene expression data used in the three biological analyses.

We used seed 42 in our n2v2r python code (*random*.*seed, numpy*.*random*.*seed*), controlling the simulated network generation and the *prerank* function of GSEApy. We found that the *prerank* function does not produce consistent GSEA plots despite using a fixed seed, and different runs can produce slightly different *p*-values. This can affect the order of the top pathways in bubble plots and can cause pathways to move into or out of the top 30, particularly when they have near identical adjusted *p*-values. Gene expression preprocessing and network inference for both bulk and single-cell data follow standard protocols as provided by the respective tools. We used default parameters in all analyses, except for single-cell metacell creation, as we needed to ensure a sufficient number of metacells for hdWGCNA. For soft-thresholding in WGCNA we used 10 as it was the middle value in the [1, 20] range used by the function *pickSoftThreshold*. For soft-thresholding using hdWGCNA we used 5 as it was the most common value between all networks for function *TestSoftPowers*.

## 7 Acknowledgments

The authors would like to thank Camila Lopes-Ramos for insightful discussions regarding sex differences. This work was supported by grants from the National Institutes of Health: P.M., V.F., C.C., J.F., E.S., and J.Q. were supported by R35CA220523; J.Q. was also supported by U24CA231846, P50CA127003, R01HG011393; E.S. was also supported by the American Lung Association grant LCD-821824. I.G. was supported by the Additional Funding Programme for Mathematical Sciences, delivered by EPSRC (EP/V521917/1) and the Heilbronn Institute for Mathematical Research.

## Supplementary Material

**Figure S1.**
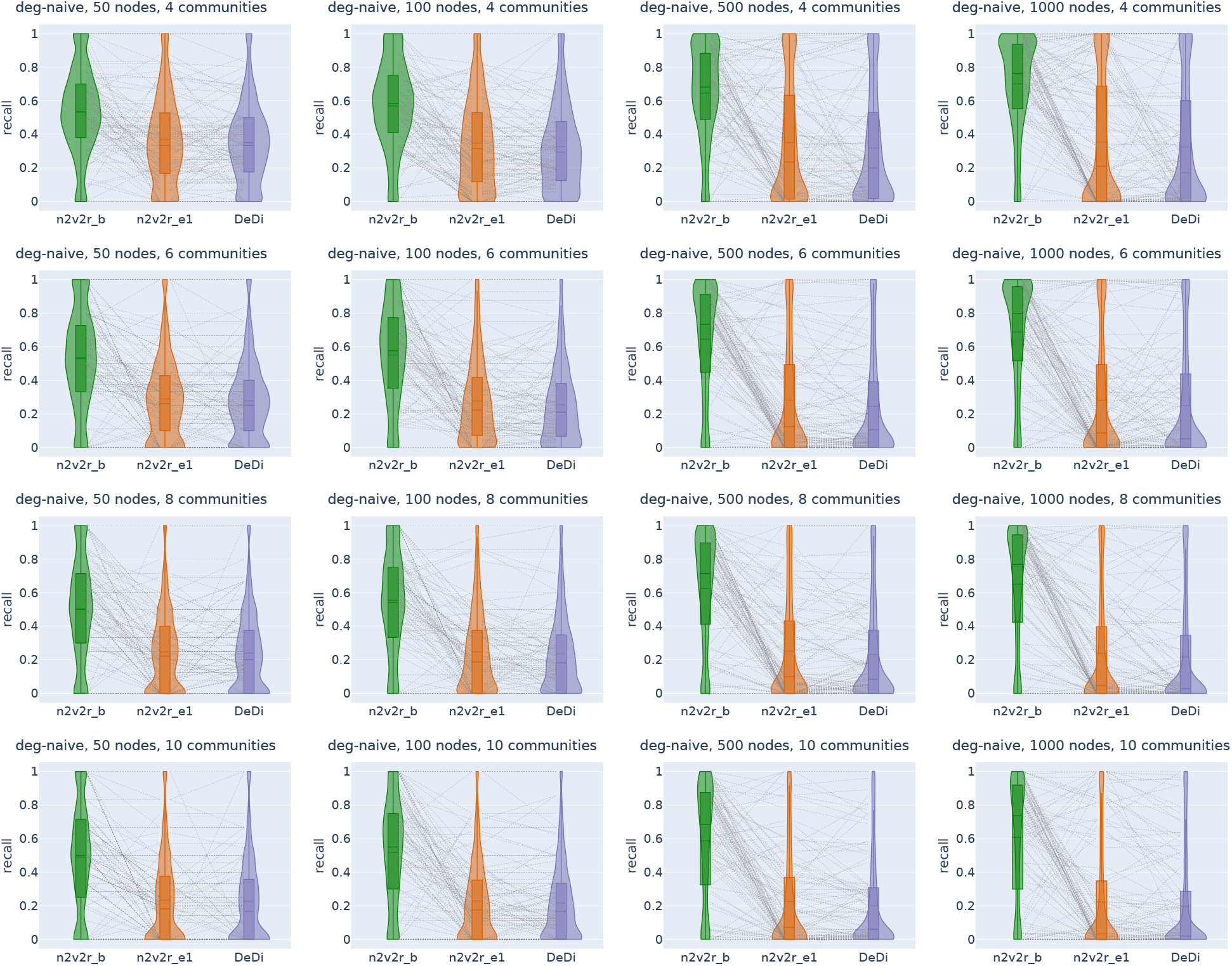
Recall violin plots for retrieving the nodes that have undergone the most significant changes between two graphs with degree-naive differences. We intervene by permuting the community probabilities of a single community to simulate changes in connections, but not in degree. We generated 2000 pairs of graphs by randomly applying this intervention with Gaussian noise. The goal is to retrieve the most changing nodes of the intervened community. The number of nodes increases across columns, while the number of communities increases down the rows. For a random subset of 100(5%) sampled graph pairs, we connected the corresponding method recall estimates. The solid and dotted lines in the violins correspond to the median and the mean, respectively.

**Figure S2.**
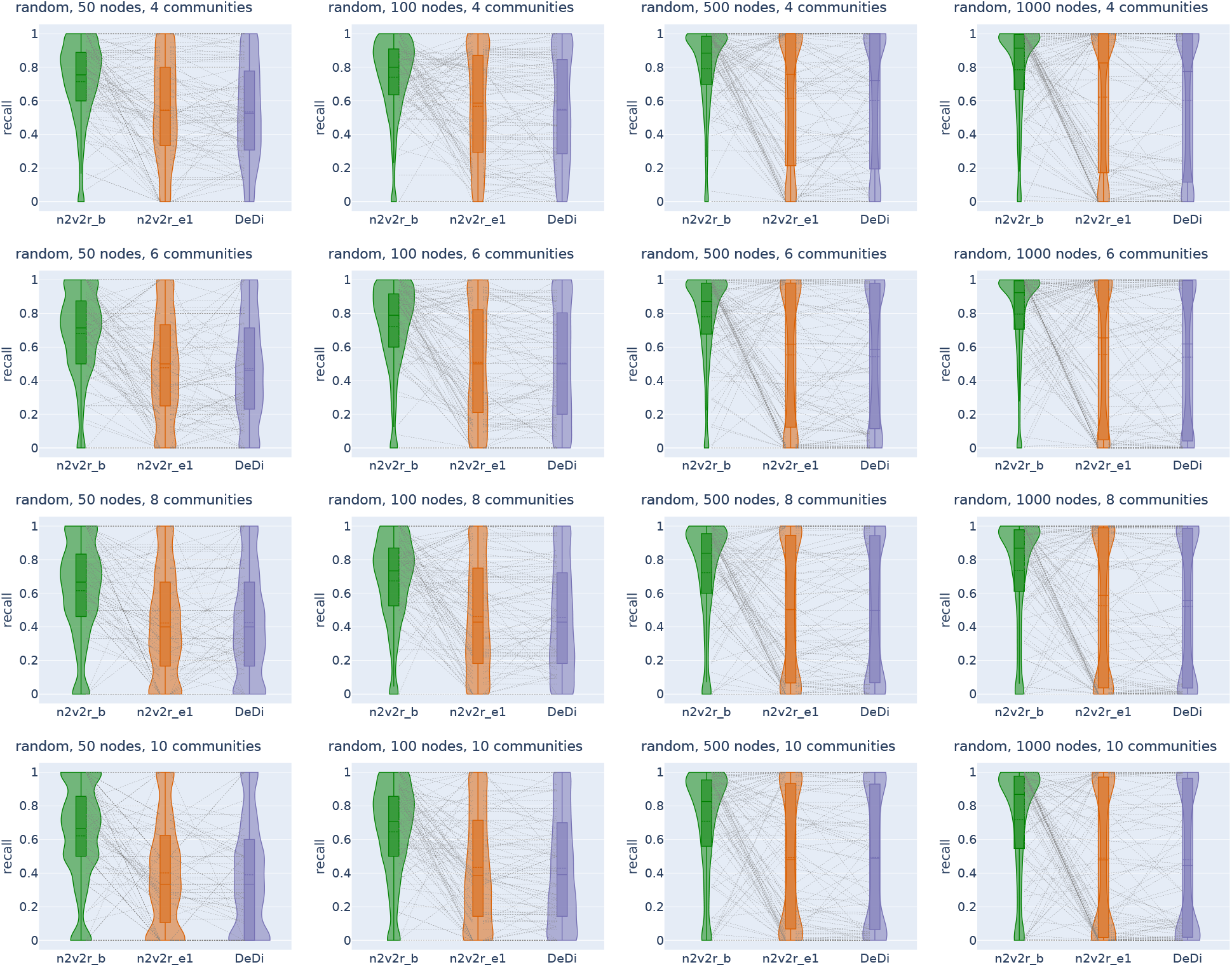
Recall violin plots for retrieving the most changing nodes between two graphs with random differences. We intervene by randomly changing the community probabilities of a single community and then updating the second adjacency matrix accordingly. We generated 2000 pairs of graphs by randomly applying this intervention with Gaussian noise. The goal is to retrieve the most changing nodes of the intervened community. The number of nodes increases across columns, while the number of communities increases down the rows. For a random subset of 100(5%) sampled graph pairs, we connect the corresponding method recall estimates. The solid and dotted lines in the violins correspond to the median and the mean, respectively.

**Figure S3.**
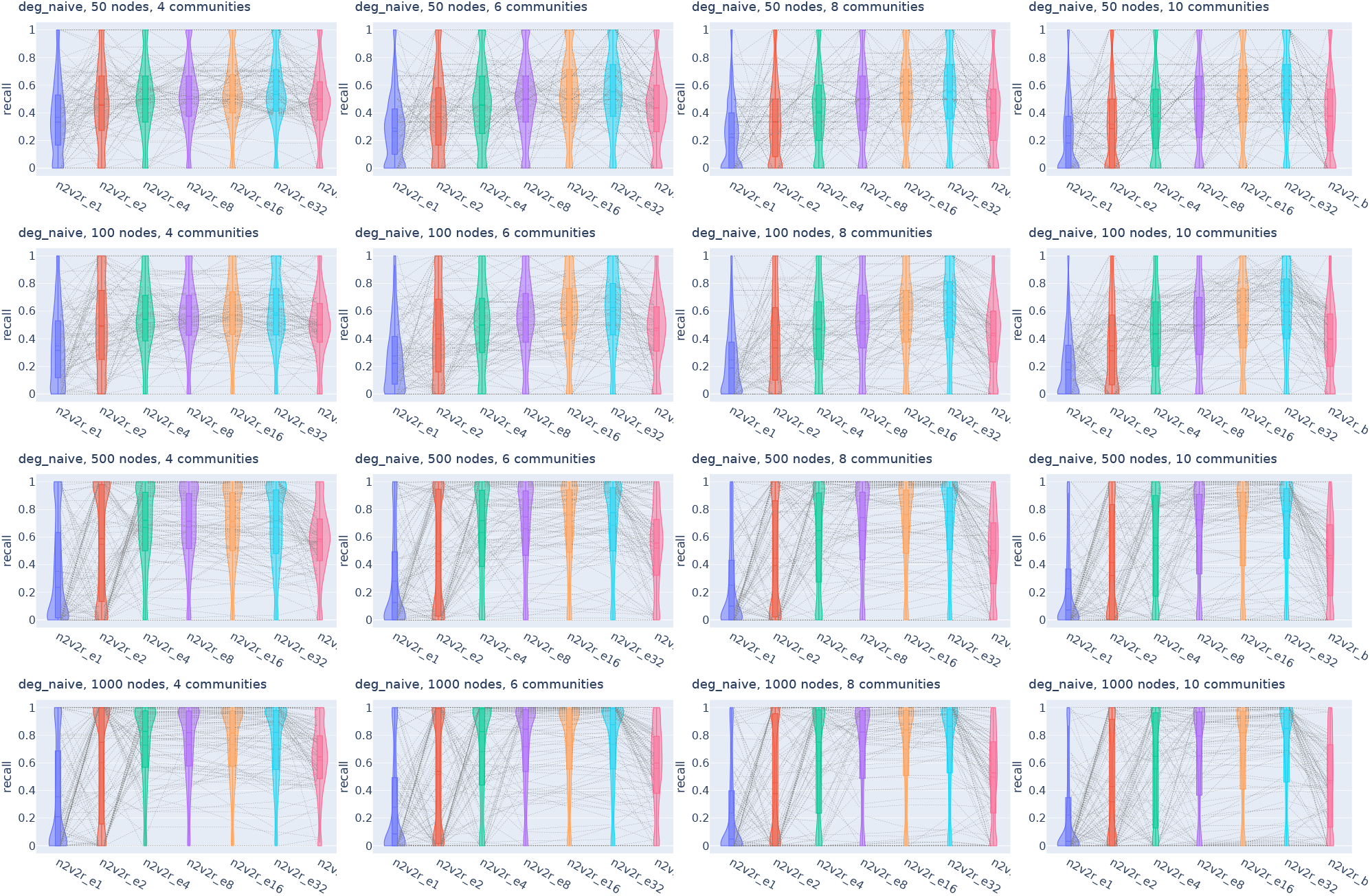
Violin plots for recall in retrieving the most changing nodes between two graphs with degree-naive differences for Euclidean distance and increasing embedding dimensionality and Borda aggregation. We intervene by permuting the community probabilities of a single community to simulate changes in connections, but not in degree. We generated 2000 pairs of graphs by randomly applying this intervention with Gaussian noise. The goal is to retrieve the most changing nodes of the intervened community. For a random subset of 100(5%) sampled graph pairs, we connect the corresponding method recall estimates. The solid and dotted lines in the violins correspond to the median and the mean, respectively.

**Figure S4.**
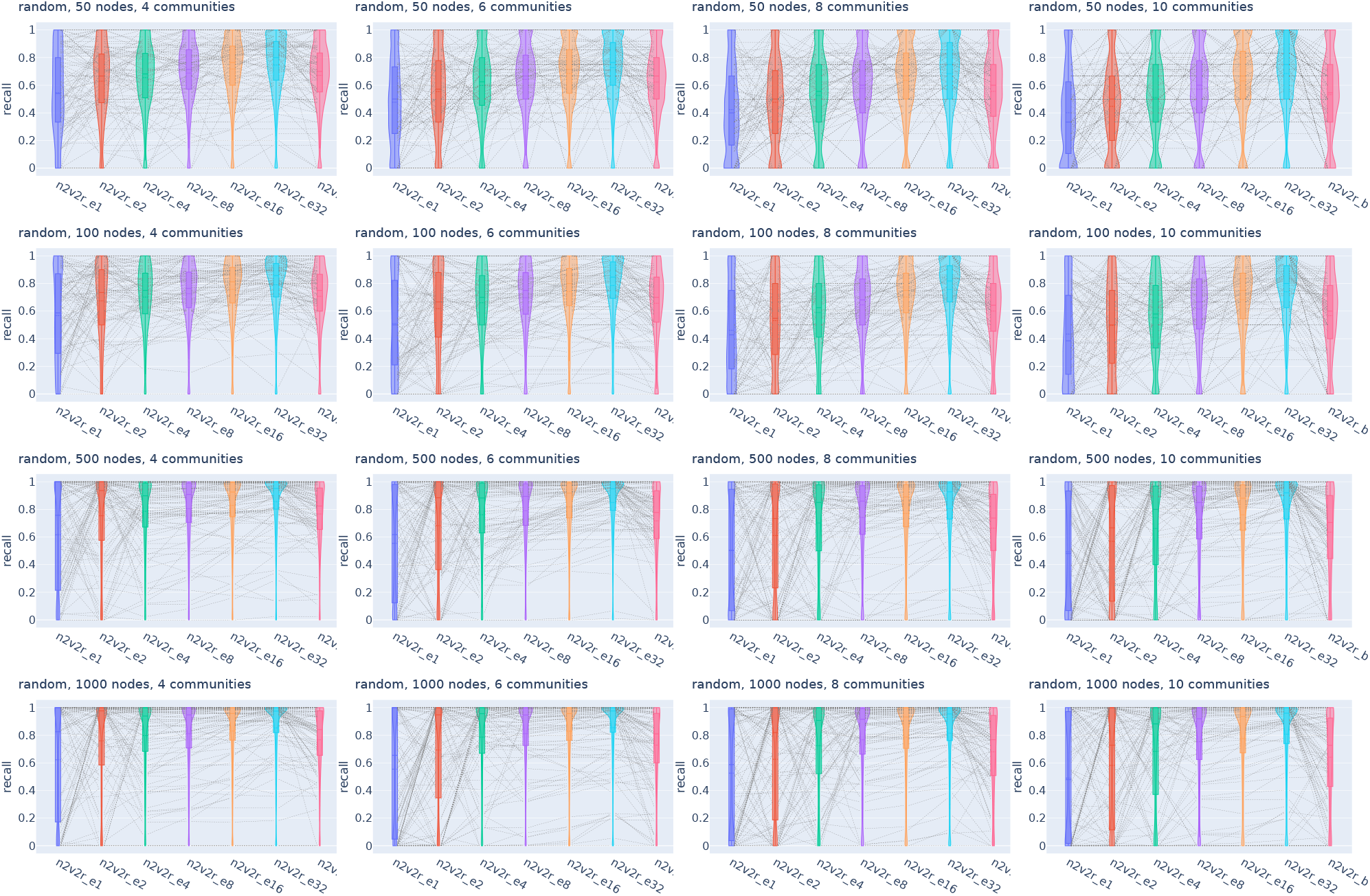
Violin plots for recall in retrieving the most changing nodes between two graphs with random differences for Euclidean distance and increasing embedding dimensionality and Borda aggregation. We intervene by randomly changing the community probabilities of a single community. We generated 2000 pairs of graphs by randomly applying this intervention with Gaussian noise. The goal is to retrieve the most changing nodes of the intervened community. For a random subset of 100(5%) sampled graph pairs, we connect the corresponding method recall estimates. The solid and dotted lines in the violins correspond to the median and the mean, respectively.

**Figure S5.**
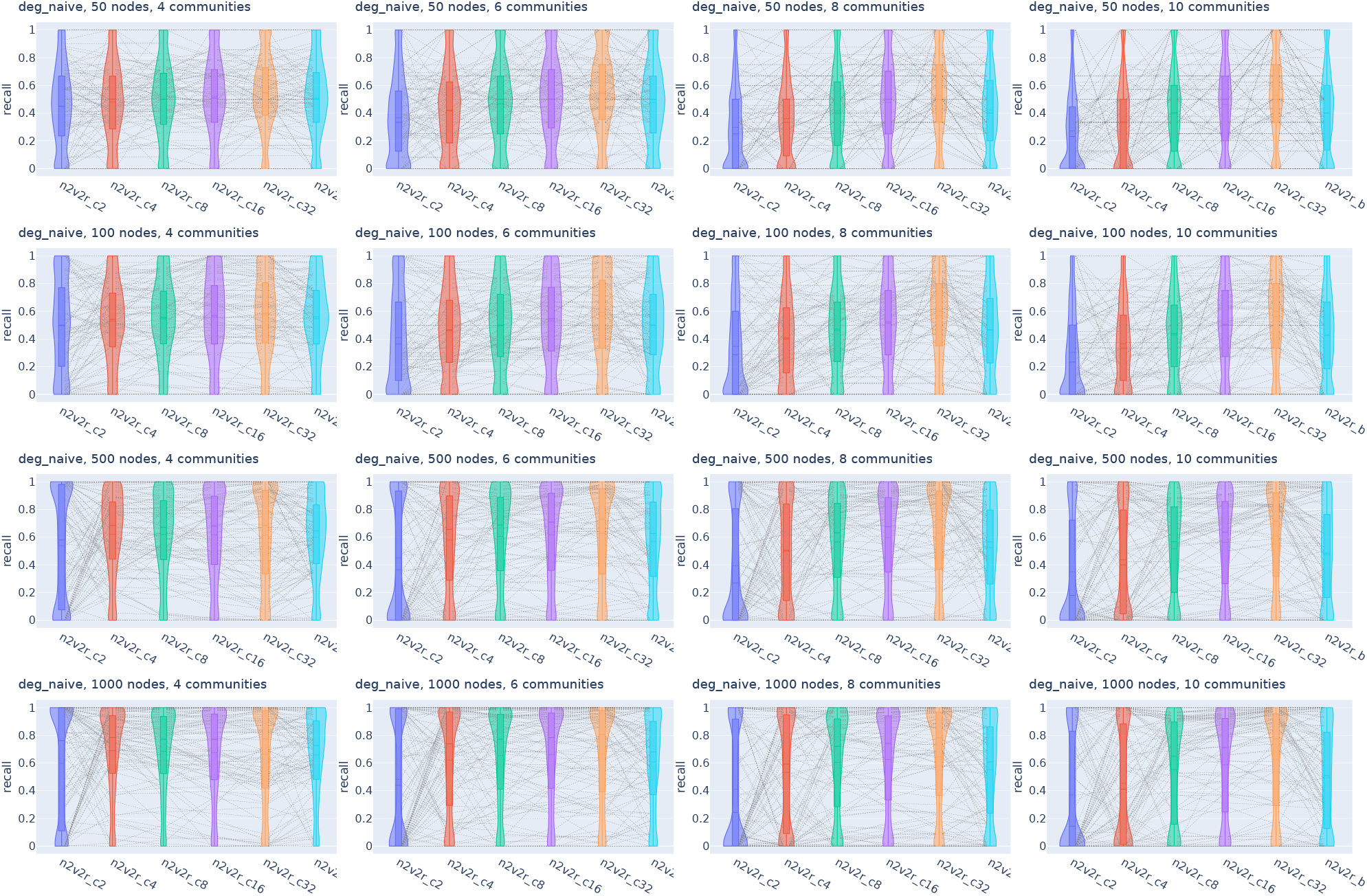
Violin plots for recall in retrieving the most changing nodes between two graphs with degree-naive differences for cosine distance and increasing embedding dimensionality and Borda aggregation. We intervene by permuting the community probabilities of a single community to simulate changes in connections, but not in degree. We generated 2000 pairs of graphs by randomly applying this intervention with Gaussian noise. The goal is to retrieve the most changing nodes of the intervened community. For a random subset of 100(5%) sampled graph pairs, we connect the corresponding method recall estimates. The solid and dotted lines in the violins correspond to the median and the mean, respectively.

**Figure S6.**
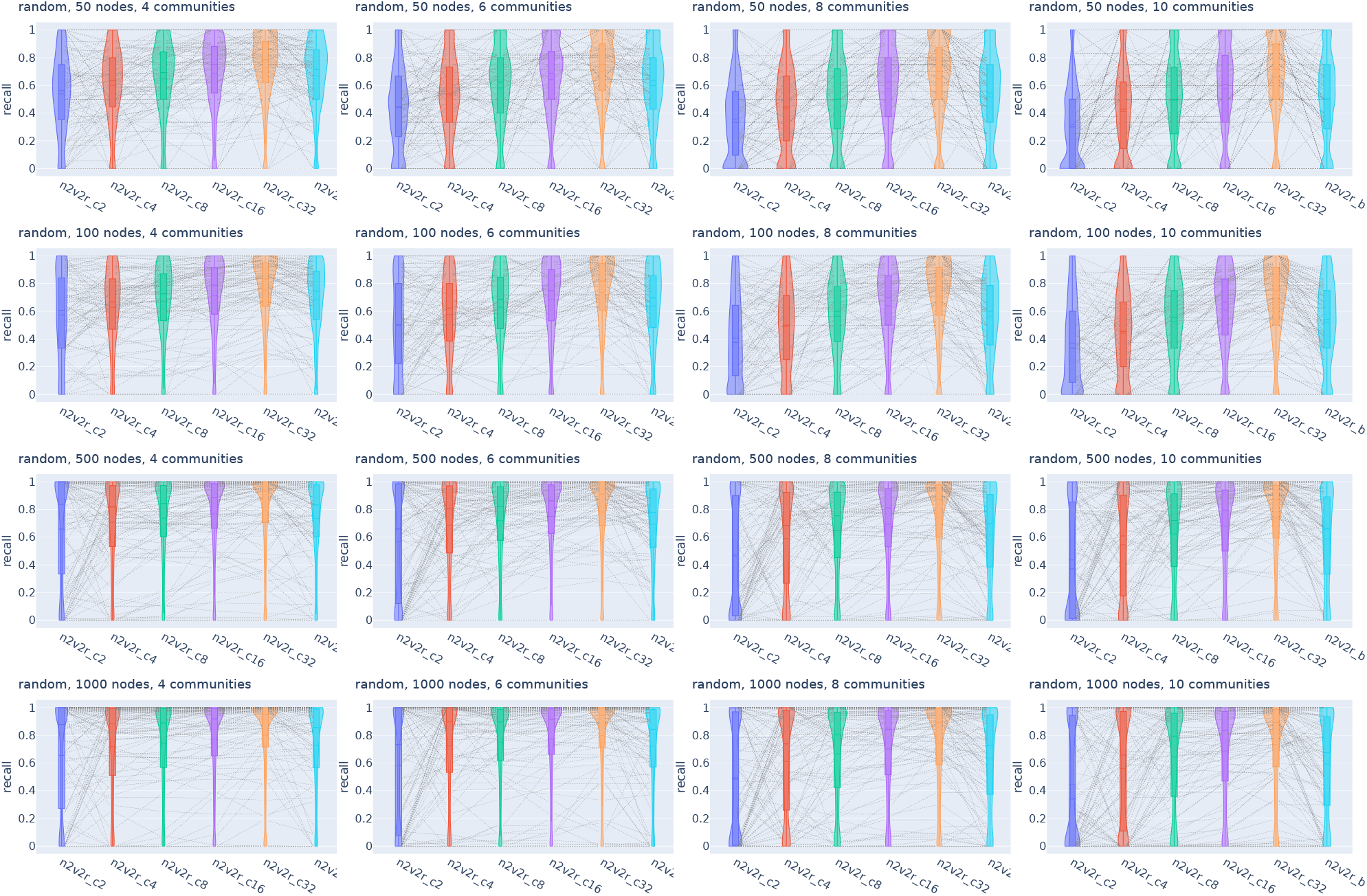
Violin plots for recall in retrieving the most changing nodes between two graphs with random differences for cosine distance and increasing embedding dimensionality and Borda aggregation. We intervene by randomly changing the community probabilities of a single community. We generate 2000 pairs of graphs by randomly applying this intervention with Gaussian noise. The goal is to retrieve the most changing nodes of the intervened community. For a random subset of 100(5%) sampled graph pairs, we connect the corresponding method recall estimates. The solid and dotted lines in the violins correspond to the median and the mean, respectively.

**Figure S7.**
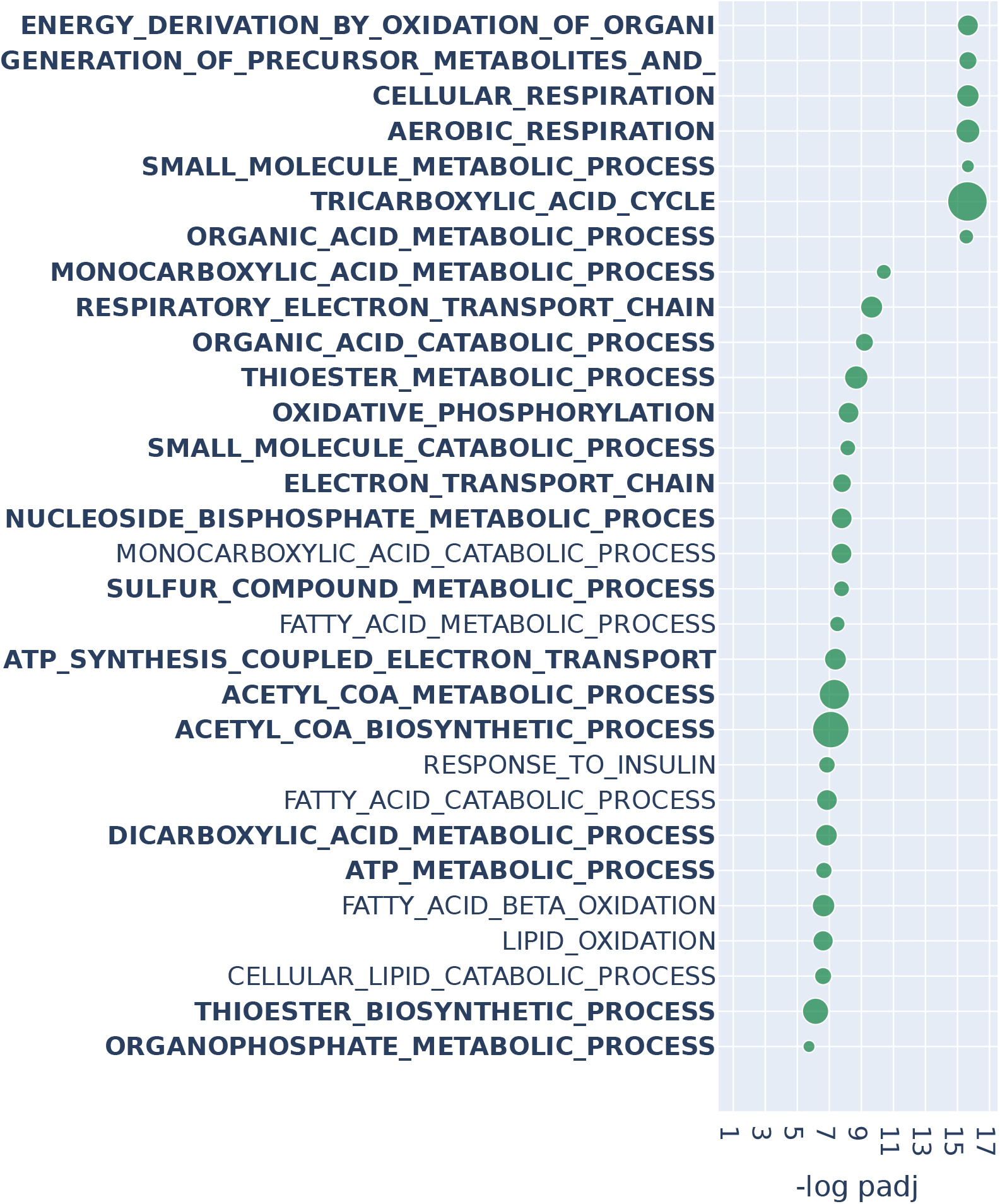
Over-representation analysis with GOBP (Gene Ontology, Biological Process) annotations on BRCA TCGA using DeDi. We consider the top 2% of genes (roughly 500) of DeDi on BRCA TCGA data when comparing luminal A tumor samples against matched adjacent normal samples using PANDA regulatory networks. We present the top 30 results ranked by adjusted p-values that pass a 0.1 FDR cutoff (equivalent to − log padj = 1). Bold indicates pathways common with n2v2r when performing the same analysis. The size of the points indicates the degree of overlap between the leading gene sets and the pathways. Long pathway names have been truncated.

**Figure S8.**
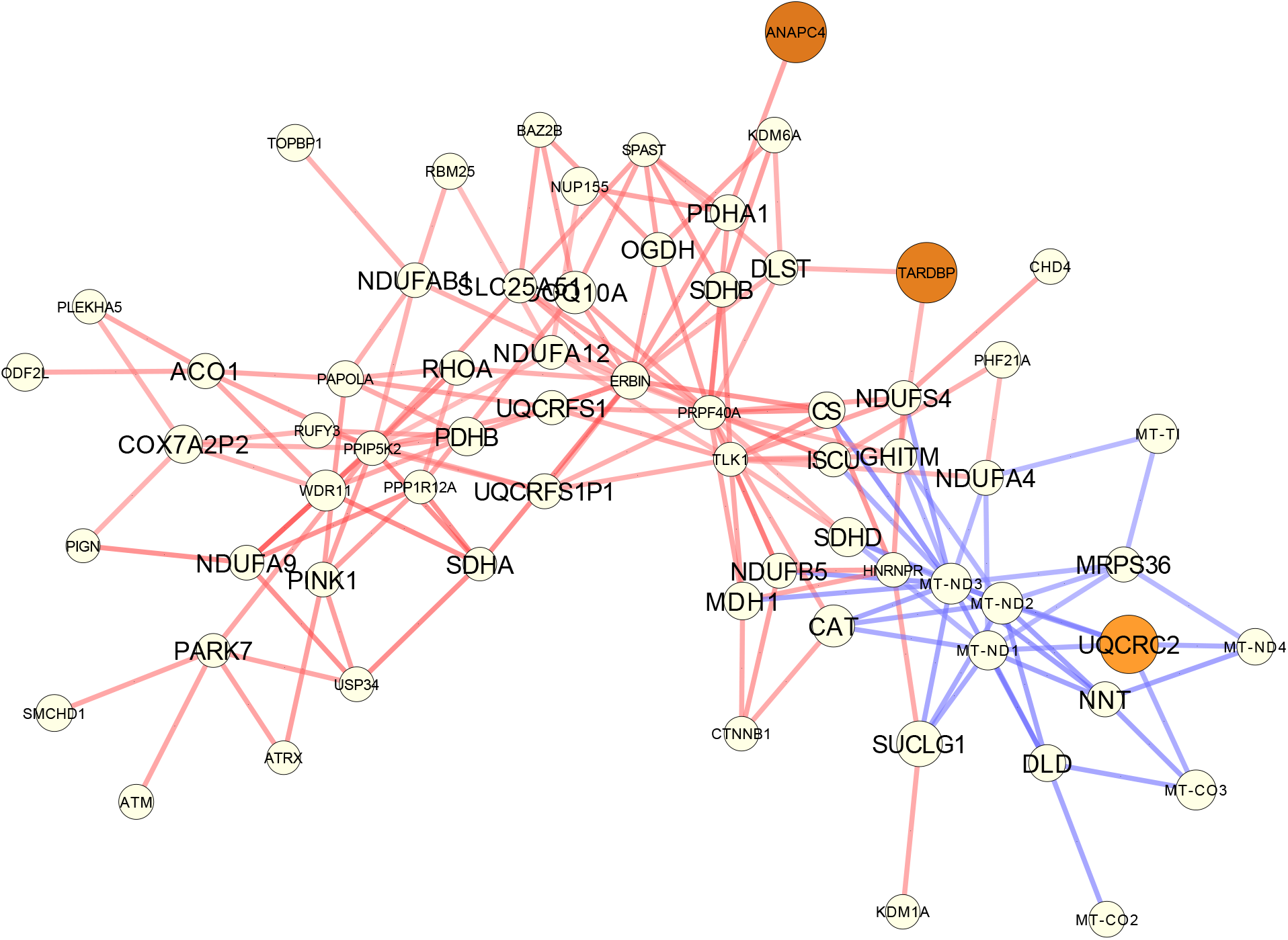
The subgraph corresponding to the differential unipartite PANDA network *A*^lumA^–*A*^normal^ focusing on the leading genes of n2v2r on the cellular respiration GOBP pathway. For every such leading gene (shown here with larger fonts), we show its top 5 neighbors (in absolute edge weight). Red edges mean higher co-regulation in luminal A patients and blue edges in their adjacent normal samples. The size of the nodes is the absolute degree difference. Nodes are colored according to the FDR-adjusted p-value from differential gene expression analysis with DESeq2: white is not significant, yellow to red is significant between 0.1 and 0.

**Figure S9.**
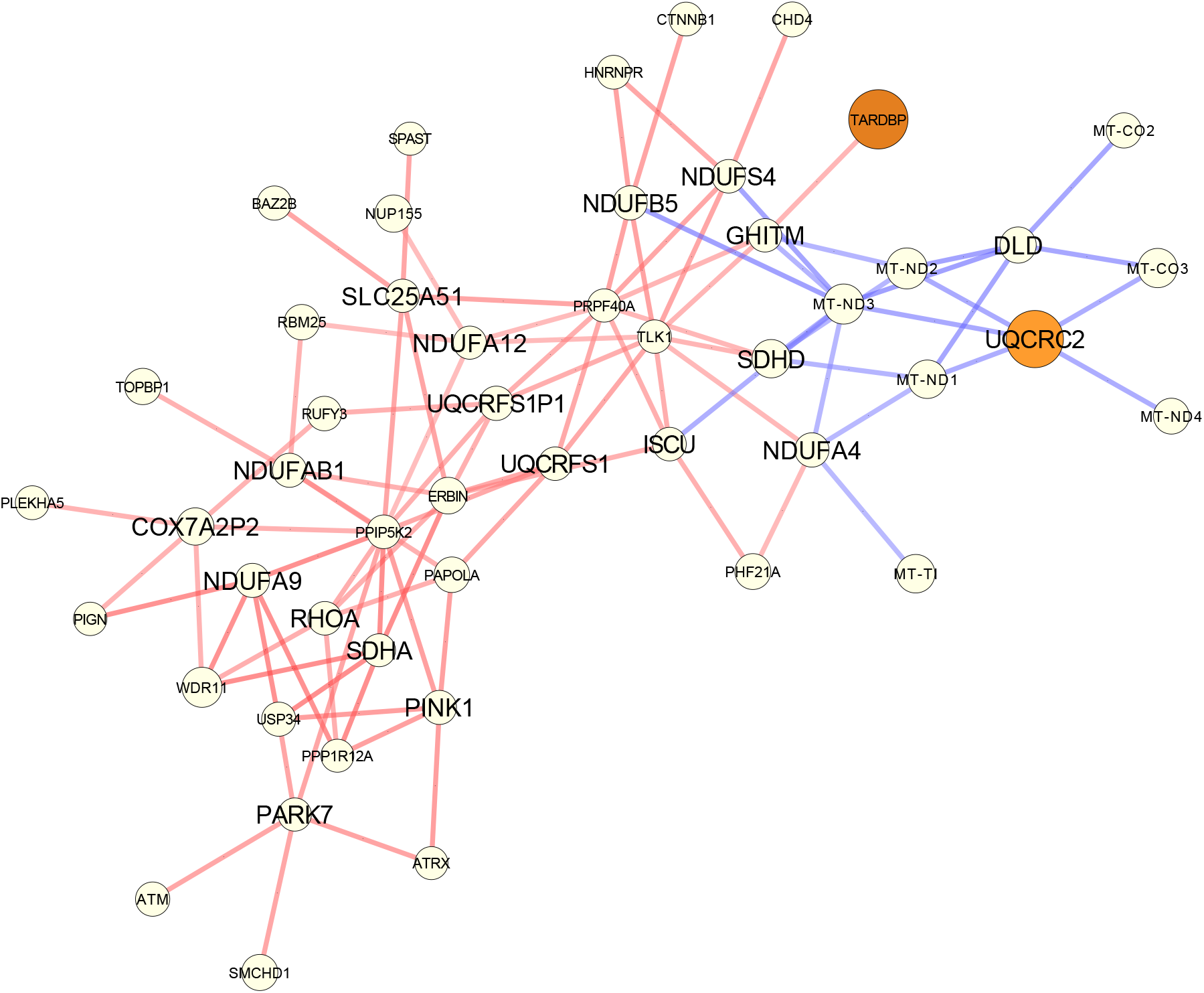
The subgraph corresponding to the differential unipartite PANDA network *A*^lumA^–*A*^normal^ focusing on the leading genes of n2v2r on the oxidative phosphorylation GOBP pathway. For every such leading gene (shown here with larger fonts), we show their top 5 neighbors (in absolute edge weight). Red edges indicate higher co-regulation in luminal A patients, and blue edges indicate higher co-regulation in their adjacent normal samples. The size of the nodes is the absolute degree difference. Nodes are colored according to the FDR-adjusted p-value from differential gene expression analysis with DESeq2: white is not significant, yellow to red is significant between 0.1 and 0.

**Figure S10.**
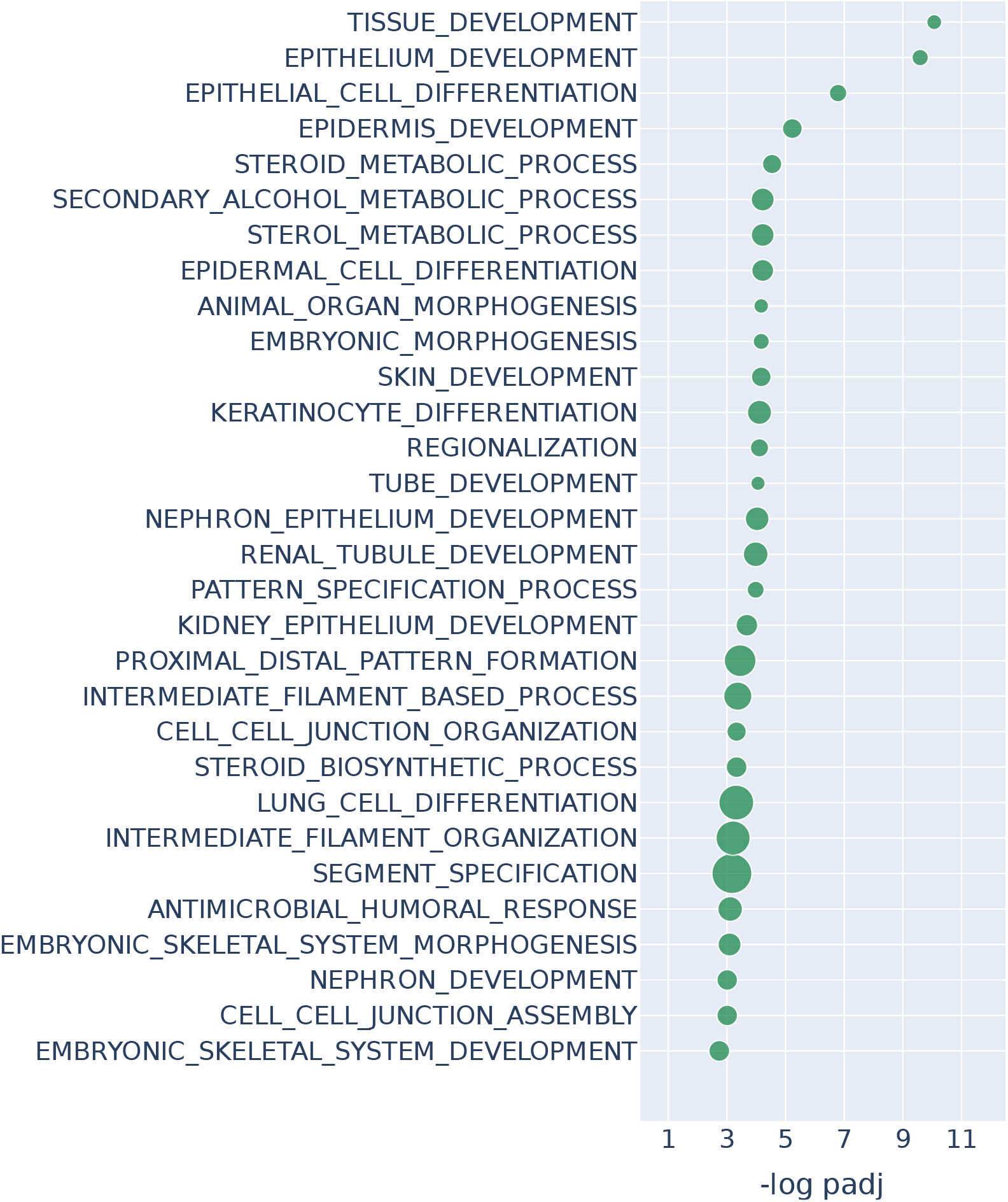
Over-representation analysis with GOBP (Gene Ontology, Biological Process) annotations on BRCA TCGA using DESeq2. We consider the top 2% of genes (roughly 500) of differential gene expression with DESeq2 on BRCA TCGA data when comparing luminal A tumor samples against matched adjacent normal samples using PANDA regulatory networks. We present the top 30 results ranked by adjusted p-values that pass a 0.1 FDR cutoff (equivalent to − log padj = 1). The size of the points indicates the degree of overlap between the leading gene sets and the pathways. Long pathway names have been truncated.

**Figure S11.**
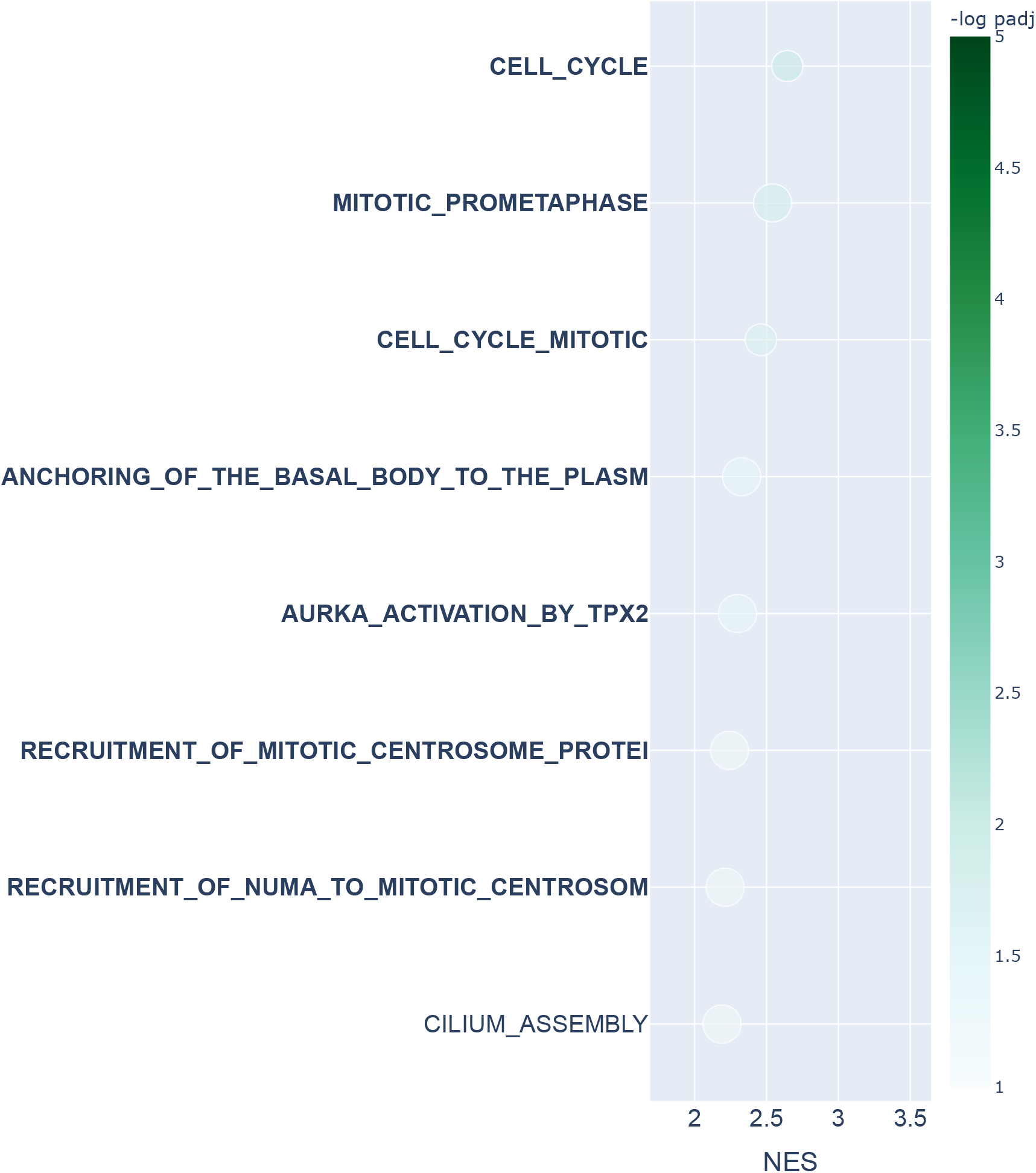
Single-cell cell-cycle transition analysis Reactome results for DeDi. The HeLa S3 cell line is integrated using Seurat, with cells aggregated to create metacells, from which four co-expression networks are generated using hdWGCNA. The four networks correspond to the cell-cycle phases G1, S, G2, and M. In this bubbleplot, gene set enrichment analysis with the Reactome database is performed on the ranked list of DeDi results to study the G1 to S transition, showing the top 30 pathways ranked by adjusted p-values that survive FDR 0.1 cutoff (equivalent to − log padj = 1). The x-axis represents the normalized enrichment score, and color the − log padj value. The size of the points indicates the degree of overlap between the leading gene sets and the pathways. Bold color indicates common pathways with n2v2r. The pathways that explicitly mention terms related to the G1 to S transition are shown in green boxes (there were none for DeDi). The same procedure for the G2 to M transition was performed, but no significant results were obtained. Long pathway names have been truncated.

**Figure S12.**
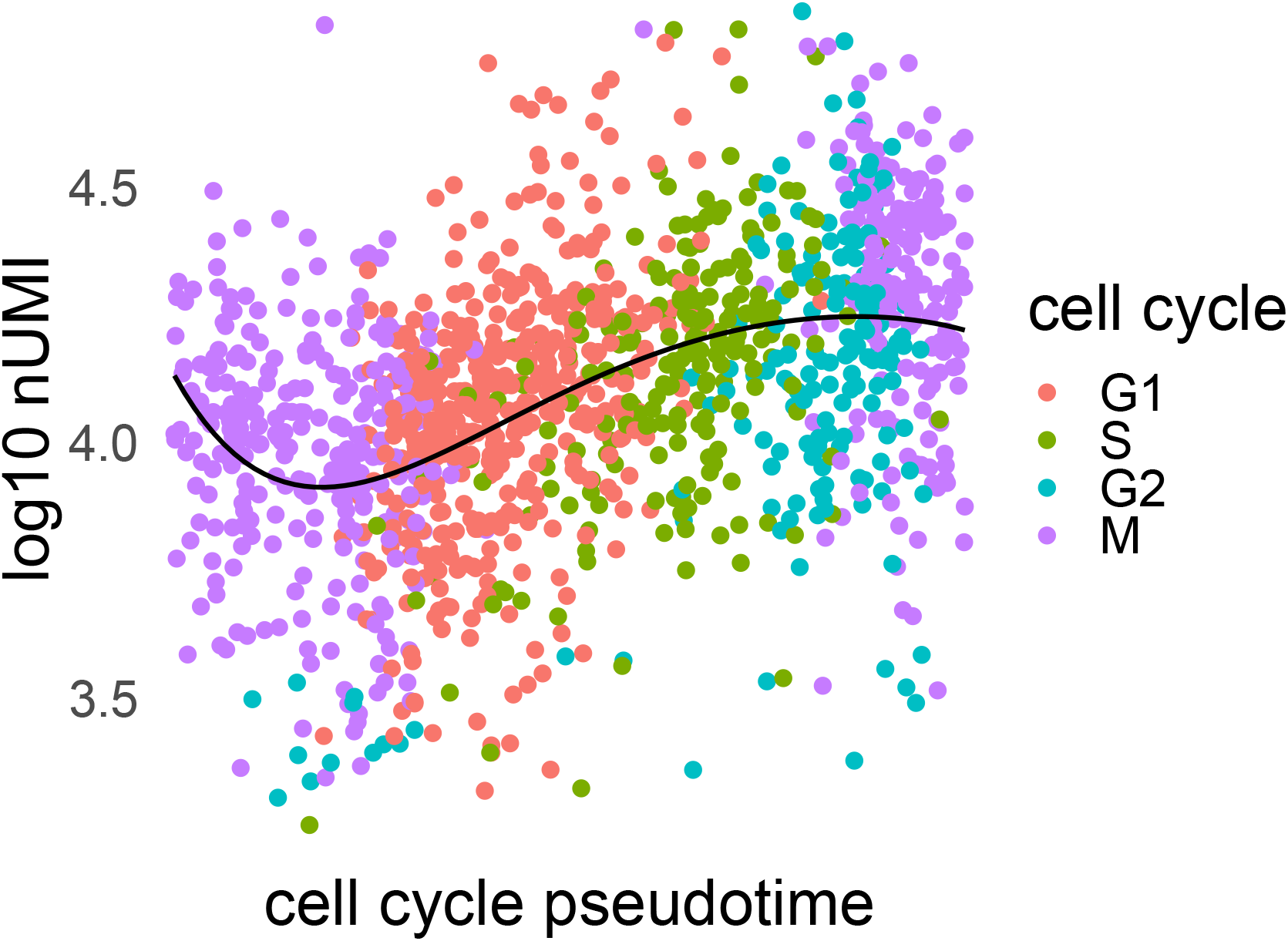
Logarithm of number of unique molecular identifiers per cell sorted according to cell cycle pseudotime and colored by cell cycle phase. The black curve is a fifth-order polynomial fit.

**Figure S13.**
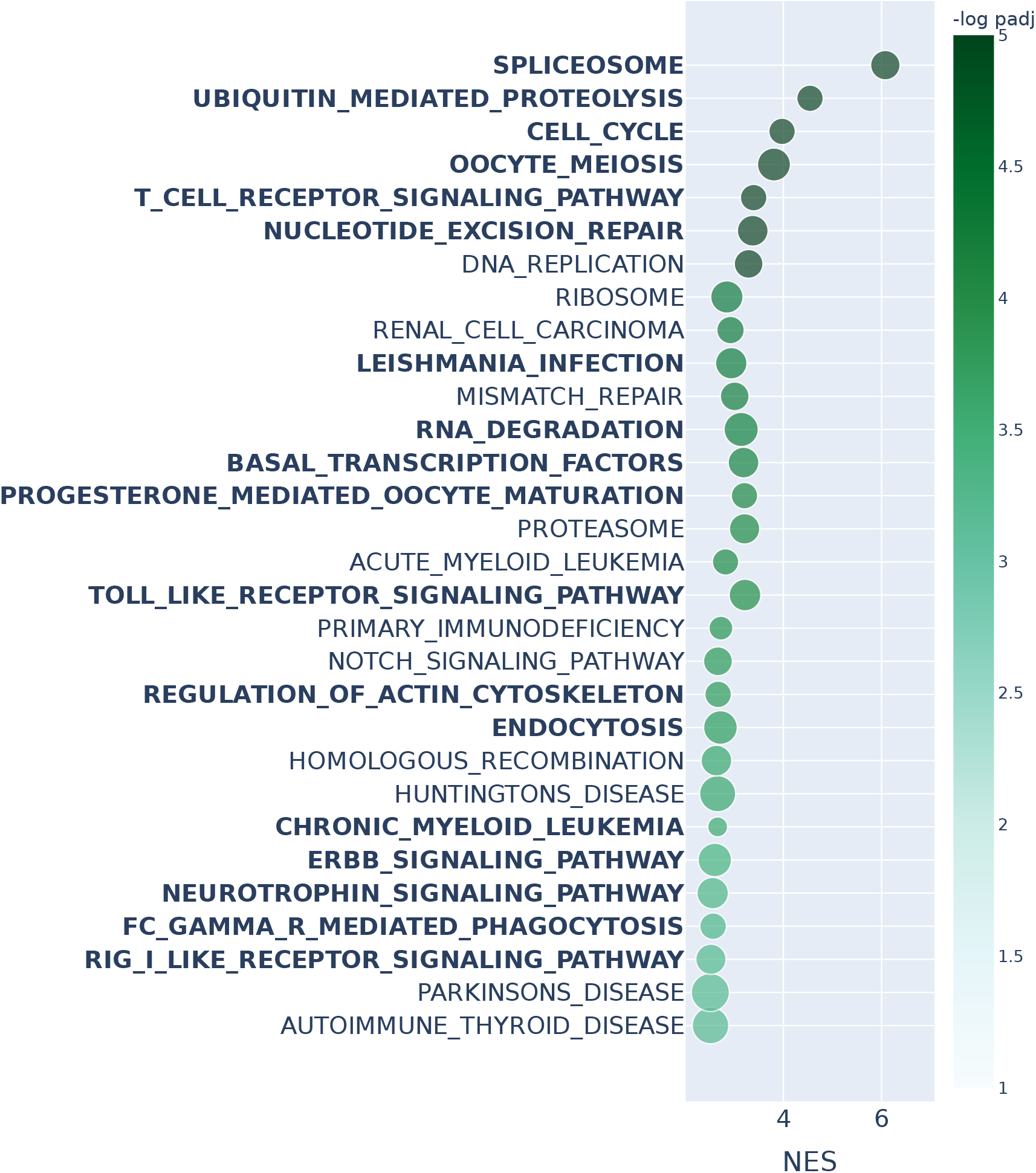
Gene set enrichment analysis with KEGG annotations and Kolmogorov–Smirnov test on TCGA LUAD using DeDi. We consider the entire DeDi ranking of sex-specific differences found in WGCNA co-expression networks inferred for TCGA LUAD male and female samples. The top 30 results passing a 0.1 FDR cutoff (equivalent to − log padj = 1) and ranked by adjusted p-values are shown. Bold indicates pathways common with n2v2r when performing the same analysis. The x-axis represents the normalized enrichment score, and the color represents the − log padj value. The size of the points indicates the degree of overlap between the leading gene sets and the pathways. Long pathway names have been truncated.

**Figure S14.**
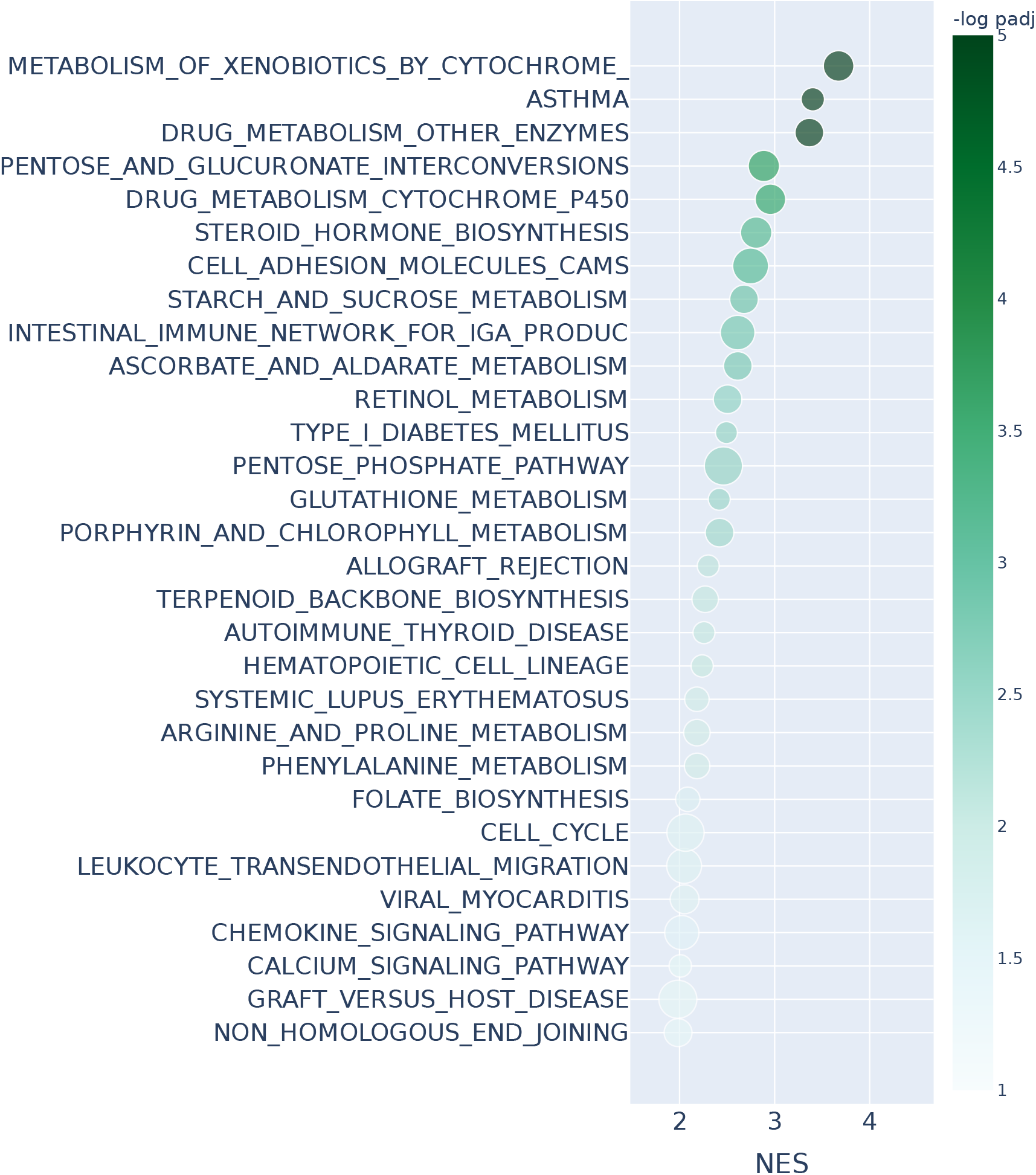
Gene set enrichment analysis with KEGG annotations and Kolmogorov–Smirnov test on TCGA LUAD using DESeq2. We consider the entire ranking from differential gene expression using DESeq2 of sex-specific differences found in WGCNA co-expression networks inferred for TCGA LUAD male and female samples. The top 30 results passing a 0.1 FDR cutoff (equivalent to − log padj = 1) and ranked by adjusted p-values are shown. The x-axis represents the normalized enrichment score, and the color represents the − log padj value. The size of the points indicates the degree of overlap between the leading gene sets and the pathways. Long pathway names have been truncated.

## References

1. D. Aran, M. Sirota, and A. J. Butte. 2015. Systematic pan-cancer analysis of tumour purity. Nature Communications 6, 1 (Dec. 2015), 8971.

2. J. Arroyo, A. Athreya, J. Cape, G. Chen, C. E. Priebe, and J. T. Vogelstein. 2021. Inference for Multiple Heterogeneous Networks with a Common Invariant Subspace. J Mach Learn Res 22, 141 (March 2021), 1–49.

3. M. Ashburner, C. A. Ball, J. A. Blake, D. Botstein, H. Butler, J. M. Cherry, A. P. Davis, K. Dolinski, S. S. Dwight, J. T. Eppig, M. A. Harris, D. P. Hill, L. Issel-Tarver, A. Kasarskis, S. Lewis, J. C. Matese, J. E. Richardson, M. Ringwald, G. M. Rubin, and G. Sherlock. 2000. Gene Ontology: tool for the unification of biology. Nat Genet 25, 1 (May 2000), 25–29.

4. A. Avagliano, M. R. Ruocco, F. Aliotta, I. Belviso, A. Accurso, S. Masone, S. Montagnani, and A. Arcucci. 2019. Mitochondrial Flexibility of Breast Cancers: A Growth Advantage and a Therapeutic Opportunity. Cells 8, 5 (Apr 2019).

5. M. Ben Guebila, T. Wang, C. M. Lopes-Ramos, V. Fanfani, D. Weighill, R. Burkholz, D. Schlauch, J. N. Paulson, M. Altenbuchinger, K. H. Shutta, A. R. Sonawane, J. Lim, G. Calderer, D. G. P. van IJzendoorn, D. Morgan, A. Marin, C.-Y. Chen, Q. Song, E. Saha, D. L. DeMeo, M. Padi, J. Platig, M. L. Kuijjer, K. Glass, and J. Quackenbush. 2023. The Network Zoo: a multilingual package for the inference and analysis of gene regulatory networks. Genome Biology 24, 1 (March 2023), 45.

6. Cancer Genome Atlas Research Network, J. N. Weinstein, E. A. Collisson, G. B. Mills, K. R. Shaw, B. A. Ozenberger, K. Ellrott, I. Shmulevich, C. Sander, and J. M. Stuart. 2013. The Cancer Genome Atlas Pan-Cancer analysis project. Nat Genet 45, 10 (Oct. 2013), 1113–1120. DOI:10.1038/ng.2764

7. V. Cappelletti, E. Iorio, P. Miodini, M. Silvestri, M. Dugo, and M. G. Daidone. 2017. Metabolic footprints and molecular subtypes in breast cancer. Dis. Markers 2017 (Dec. 2017), 7687851.

8. N. Dingwall and C. Potts. 2018. Mittens: an Extension of GloVe for Learning Domain-Specialized Representations. In Proceedings of the 2018 Conference of the North American Chapter of the Association for Computational Linguistics: Human Language Technologies, Volume 2 (Short Papers), Marilyn Walker, Heng Ji, and Amanda Stent (Eds.). Association for Computational Linguistics, New Orleans, Louisiana, 212–217. DOI:10.18653/v1/N18-2034

9. R. El-Botty, L. Morriset, E. Montaudon, Z. Tariq, A. Schnitzler, M. Bacci, N. Lorito, L. Sourd, L. Huguet, A. Dahmani, P. Painsec, H. Derrien, S. Vacher, J. Masliah-Planchon, V. Raynal, S. Baulande, T. Larcher, A. Vincent-Salomon, G. Dutertre, P. Cottu, G. Gentric, F. Mechta-Grigoriou, S. Hutton, K. Driouch, I. Bièche, A. Morandi, and E. Marangoni. 2023. Oxidative phosphorylation is a metabolic vulnerability of endocrine therapy and palbociclib resistant metastatic breast cancers. Nature Communications 14, 1 (14 Jul 2023), 4221. DOI:10.1038/s41467-023-40022-5

10. T. Fan, S. Pan, S. Yang, B. Hao, L. Zhang, D. Li, and Q. Geng. 2021. Clinical Significance and Immunologic Landscape of a Five-IL(R)-Based Signature in Lung Adenocarcinoma. Front Immunol 12 (Aug. 2021), 693062.

11. Z. Fang, X. Liu, and G. Peltz. 2022. GSEApy: a comprehensive package for performing gene set enrichment analysis in Python. Bioinformatics 39, 1 (11 2022), btac757. DOI:10.1093/bioinformatics/btac757 arXiv:https://academic.oup.com/bioinformatics/article-pdf/39/1/btac757/48448971/btac757.pdf

12. A. Frankish, M. Diekhans, I. Jungreis, J. Lagarde, J. Loveland, J. M. Mudge, C. Sisu, J. C. Wright, J. Armstrong, I. Barnes, A. Berry, A. Bignell, C. Boix, S. Carbonell Sala, F. Cunningham, T. Di Domenico, S. Donaldson, I. Fiddes, C. García Girón, J. M. Gonzalez, T. Grego, M. Hardy, T. Hourlier, K. L. Howe, T. Hunt, O. G. Izuogu, R. Johnson, F. J. Martin, L. Martínez, S. Mohanan, P. Muir, F. C. P. Navarro, A. Parker, B. Pei, F. Pozo, F. C. Riera, M. Ruffier, B. M. Schmitt, E. Stapleton, M.-M. Suner, I. Sycheva, B. Uszczynska-Ratajczak, M. Y. Wolf, J. Xu, Y. Yang, A. Yates, D. Zerbino, Y. Zhang, J. Choudhary, M. Gerstein, R. Guigó, T. J. P. Hubbard, M. Kellis, B. Paten, M. L. Tress, and P. Flicek. 2020. GENCODE 2021. Nucleic Acids Research 49, D1 (12 2020), D916–D923. DOI:10.1093/nar/gkaa1087

13. N. Fuentes, M. S. Rodriguez, and P. Silveyra. 2021. Role of sex hormones in lung cancer. Experimental Biology and Medicine 246, 19 (2021), 2098–2110. DOI:10.1177/15353702211019697 arXiv:https://doi.org/10.1177/15353702211019697 PMID: 34080912.

14. I. Gallagher, A. Jones, A. Bertiger, C. E. Priebe, and P. Rubin-Delanchy. 2023. Spectral embedding of weighted graphs. J. Amer. Statist. Assoc. (2023), 1–10.

15. I. Gallagher, A. Jones, and P. Rubin-Delanchy. 2021. Spectral embedding for dynamic networks with stability guarantees. Advances in Neural Information Processing Systems 34 (2021), 10158–10170.

16. Y. Gautam, Y. Afanador, T. Abebe, J. E. López, and T. B. Mersha. 2019. Genome-wide analysis revealed sex-specific gene expression in asthmatics. Hum Mol Genet 28, 15 (Aug. 2019), 2600–2614.

17. M. Gillespie, B. Jassal, R. Stephan, M. Milacic, K. Rothfels, A. Senff-Ribeiro, J. Griss, C. Sevilla, L. Matthews, C. Gong, C. Deng, T. Varusai, E. Ragueneau, Y. Haider, B. May, V. Shamovsky, J. Weiser, T. Brunson, N. Sanati, L. Beckman, X. Shao, A. Fabregat, K. Sidiropoulos, J. Murillo, G. Viteri, J. Cook, S. Shorser, G. Bader, E. Demir, C. Sander, R. Haw, G. Wu, L. Stein, H. Hermjakob, and P. D’Eustachio. 2021. The reactome pathway knowledgebase 2022. Nucleic Acids Research 50, D1 (11 2021), D687–D692. DOI:10.1093/nar/gkab1028 arXiv:https://academic.oup.com/nar/article-pdf/50/D1/D687/42058295/gkab1028.pdf

18. K. Glass, C. Huttenhower, J. Quackenbush, and G.-C. Yuan. 2013. Passing Messages between Biological Networks to Refine Predicted Interactions. PLOS ONE 8, 5 (05 2013), 1–14. DOI:10.1371/journal.pone.0064832

19. C. E. Grant, T. L. Bailey, and W. S. Noble. 2011. FIMO: scanning for occurrences of a given motif. Bioinformatics 27, 7 (Apr 2011), 1017–1018.

20. A. Grover and J. Leskovec. 2016. Node2vec: Scalable Feature Learning for Networks. In Proceedings of the 22nd ACM SIGKDD International Conference on Knowledge Discovery and Data Mining (KDD ‘16). Association for Computing Machinery, New York, NY, USA, 855–864. DOI:10.1145/2939672.2939754

21. W. L. Hamilton, R. Ying, and J. Leskovec. 2017. Inductive Representation Learning on Large Graphs. In Proceedings of the 31st International Conference on Neural Information Processing Systems (NIPS’17). Curran Associates Inc., Red Hook, NY, USA, 1025–1035.

22. Y. Hao, S. Hao, E. Andersen-Nissen, W. M. M. III, S. Zheng, A. Butler, M. J. Lee, A. J. Wilk, C. Darby, M. Zagar, P. Hoffman, M. Stoeckius, E. Papalexi, E. P. Mimitou, J. Jain, A. Srivastava, T. Stuart, L. B. Fleming, B. Yeung, A. J. Rogers, J. M. McElrath, C. A. Blish, R. Gottardo, P. Smibert, and R. Satija. 2021. Integrated analysis of multimodal single-cell data. Cell (2021). DOI:10.1016/j.cell.2021.04.048

23. P. D. Hoff, A. E. Raftery, and M. S. Handcock. 2002. Latent Space Approaches to Social Network Analysis. J. Amer. Statist. Assoc. 97, 460 (Dec. 2002), 1090–1098.

24. X. Hu, J. Li, M. Fu, X. Zhao, and W. Wang. 2021. The JAK/STAT signaling pathway: from bench to clinic. Signal Transduction and Targeted Therapy 6, 1 (Nov. 2021), 402.

25. A. Jones and P. Rubin-Delanchy. 2020. The multilayer random dot product graph. arXiv preprint 2007.10455 (2020).

26. M. Kanehisa and S. Goto. 2000. KEGG: Kyoto Encyclopedia of Genes and Genomes. Nucleic Acids Research 28, 1 (01 2000), 27–30. arXiv:https://academic.oup.com/nar/article-pdf/28/1/27/9895154/280027.pdf

27. S. Kim, D. H. Kim, W.-H. Jung, and J. S. Koo. 2013. Succinate dehydrogenase expression in breast cancer. Springerplus 2, 1 (July 2013), 299.

28. M. Kivelä, A. Arenas, M. Barthelemy, J. P. Gleeson, Y. Moreno, and M. A. Porter. 2014. Multilayer networks. Journal of Complex Networks 2, 3 (07 2014), 203–271. DOI:10.1093/comnet/cnu016 arXiv:https://academic.oup.com/comnet/article-pdf/2/3/203/9130906/cnu016.pdf

29. S. L. Klein and K. L. Flanagan. 2016. Sex differences in immune responses. Nat Rev Immunol 16, 10 (Aug. 2016), 626–638.

30. E. C. Koc, F. C. Koc, F. Kartal, M. Tirona, and H. Koc. 2022. Role of mitochondrial translation in remodeling of energy metabolism in ER/PR(+) breast cancer. Front Oncol 12 (2022), 897207.

31. J. Krumsiek, K. Mittelstrass, K. T. Do, F. Stückler, J. Ried, J. Adamski, A. Peters, T. Illig, F. Kronenberg, N. Friedrich, M. Nauck, M. Pietzner, D. O. Mook-Kanamori, K. Suhre, C. Gieger, H. Grallert, F. J. Theis, and G. Kastenmüller. 2015. Gender-specific pathway differences in the human serum metabolome. Metabolomics 11, 6 (Dec. 2015), 1815–1833.

32. P. Langfelder and S. Horvath. 2008. WGCNA: an R package for weighted correlation network analysis. BMC Bioinformatics 1 (2008), 559. https://bmcbioinformatics.biomedcentral.com/articles/10.1186/1471-2105-9-559

33. M. J. Legato, P. A. Johnson, and J. E. Manson. 2016. Consideration of Sex Differences in Medicine to Improve Health Care and Patient Outcomes. JAMA 316, 18 (11 2016), 1865–1866. DOI:10.1001/jama.2016.13995 arXiv:https://jamanetwork.com/journals/jama/articlepdf/2577141/jvp160127.pdf

34. K. Levin, A. Athreya, M. Tang, V. Lyzinski, Y. Park, and C. E. Priebe. 2019. A central limit theorem for an omnibus embedding of multiple random graphs and implications for multiscale network inference. (June 2019). 1705.09355 [stat].

35. L. D. Li, H. F. Sun, X. X. Liu, S. P. Gao, H. L. Jiang, X. Hu, and W. Jin. 2015. Down-Regulation of NDUFB9 Promotes Breast Cancer Cell Proliferation, Metastasis by Mediating Mitochondrial Metabolism. PLoS One 10, 12 (2015), e0144441.

36. X. Li, S. Wei, L. Deng, H. Tao, M. Liu, Z. Zhao, X. Du, Y. Li, and J. Hou. 2023. Sex-biased molecular differences in lung adenocarcinoma are ethnic and smoking specific. BMC Pulmonary Medicine 23, 1 (March 2023), 99.

37. C. M. Lopes-Ramos, C.-Y. Chen, M. L. Kuijjer, J. N. Paulson, A. R. Sonawane, M. Fagny, J. Platig, K. Glass, J. Quackenbush, and D. L. DeMeo. 2020. Sex Differences in Gene Expression and Regulatory Networks across 29 Human Tissues. Cell Rep 31, 12 (June 2020), 107795.

38. C. M. Lopes-Ramos, M. L. Kuijjer, S. Ogino, C. S. Fuchs, D. L. DeMeo, K. Glass, and J. Quackenbush. 2018. Gene regulatory network analysis identifies sex-linked differences in colon cancer drug metabolism. Cancer Res. 78, 19 (Oct. 2018), 5538–5547.

39. M. I. Love, W. Huber, and S. Anders. 2014. Moderated estimation of fold change and dispersion for RNA-seq data with DESeq2. Genome Biology 15, 12 (Dec. 2014), 550.

40. P. Lunetti, M. Di Giacomo, D. Vergara, S. De Domenico, M. Maffia, V. Zara, L. Capobianco, and A. Ferramosca. 2019. Metabolic reprogramming in breast cancer results in distinct mitochondrial bioenergetics between luminal and basal subtypes. FEBS J 286, 4 (Feb. 2019), 688–709.

41. A. Modell, I. Gallagher, J. Cape, and P. Rubin-Delanchy. 2022. Spectral embedding and the latent geometry of multipartite networks. (2022). arXiv:stat.ME/2202.03945

42. S. Morabito, F. Reese, N. Rahimzadeh, E. Miyoshi, and V. Swarup. 2023. hdWGCNA identifies co-expression networks in high-dimensional transcriptomics data. Cell reports Methods (2023). https://www.biorxiv.org/content/10.1101/2022.09.22.509094v1

43. J. Ortega-Lozano, L. Gómez-Caudillo, A. Briones-Herrera, O. E. Aparicio-Trejo, and J. Pedraza-Chaverri. 2022. Characterization of Mitochondrial Proteome and Function in Luminal A and Basal-like Breast Cancer Subtypes Reveals Alteration in Mitochondrial Dynamics and Bioenergetics Relevant to Their Diagnosis. Biomolecules 12, 3 (Feb. 2022).

44. C. Özdemir, C. Csajka, G.-P. Dotto, and A. D. Wagner. 2018. Sex Differences in Efficacy and Toxicity of Systemic Treatments: An Undervalued Issue in the Era of Precision Oncology. Journal of Clinical Oncology 36, 26 (2018), 2680–2683. DOI:10.1200/JCO.2018.78.3290 arXiv:https://doi.org/10.1200/JCO.2018.78.3290 PMID: 30004815.

45. L. Peel, D. B. Larremore, and A. Clauset. 2017. The ground truth about metadata and community detection in networks. Science Advances 3, 5 (May 2017), e1602548.

46. B. Perozzi, R. Al-Rfou, and S. Skiena. 2014. DeepWalk: online learning of social representations. In Proceedings of the 20th ACM SIGKDD international conference on Knowledge discovery and data mining. ACM, New York New York USA, 701–710.

47. A. Prat, E. Pineda, B. Adamo, P. Galván, A. Fernández, L. Gaba, M. Díez, M. Viladot, A. Arance, and M. Muñoz. 2015. Clinical implications of the intrinsic molecular subtypes of breast cancer. The Breast 24 (2015), S26–S35. DOI:10.1016/j.breast.2015.07.008

48. C. E. Priebe, Y. Park, J. T. Vogelstein, J. M. Conroy, V. Lyzinski, M. Tang, A. Athreya, J. Cape, and E. Bridgeford. 2019. On a two-truths phenomenon in spectral graph clustering. Proceedings of the National Academy of Sciences 116, 13 (2019), 5995–6000.

49. K. D. Reddy and B. G. G. Oliver. 2023. Sexual dimorphism in chronic respiratory diseases. Cell & Bioscience 13, 1 (March 2023), 47. DOI:10.1186/s13578-023-00998-5

50. P. S. Reel, S. Reel, E. Pearson, E. Trucco, and E. Jefferson. 2021. Using machine learning approaches for multi-omics data analysis: A review. Biotechnol Adv 49 (March 2021), 107739.

51. M. E. Ritchie, B. Phipson, D. Wu, Y. Hu, C. W. Law, W. Shi, and G. K. Smyth. 2015. limma powers differential expression analyses for RNA-sequencing and microarray studies. Nucleic Acids Research 43, 7 (01 2015), e47.#x2013;e47. arXiv:https://academic.oup.com/nar/article-pdf/43/7/e47/7207289/gkv007.pdf

52. M. D. Robinson, D. J. McCarthy, and G. K. Smyth. 2010. edgeR: a Bioconductor package for differential expression analysis of digital gene expression data. Bioinformatics 26, 1 (Jan. 2010), 139–140.

53. J. B. Rubin. 2022. The spectrum of sex differences in cancer. Trends in Cancer 8, 4 (2022), 303–315.

54. J. B. Rubin, J. S. Lagas, L. Broestl, J. Sponagel, N. Rockwell, G. Rhee, S. F. Rosen, S. Chen, R. S. Klein, P. Imoukhuede, and J. Luo. 2020. Sex differences in cancer mechanisms. Biology of Sex Differences 11, 1 (April 2020), 17. DOI:10.1186/s13293-020-00291-x

55. P. Rubin-Delanchy, J. Cape, M. Tang, and C. E. Priebe. 2022. A Statistical Interpretation of Spectral Embedding: The Generalised Random Dot Product Graph. Journal of the Royal Statistical Society Series B: Statistical Methodology 84, 4 (Sept. 2022), 1446–1473.

56. C. Ruiz, M. Zitnik, and J. Leskovec. 2021. Identification of disease treatment mechanisms through the multiscale interactome. Nat Commun 12, 1 (March 2021), 1796.

57. J. Rutter, D. R. Winge, and J. D. Schiffman. 2010. Succinate dehydrogenase - Assembly, regulation and role in human disease. Mitochondrion 10, 4 (Jun 2010), 393–401.

58. E. Saha, M. Ben Guebila, V. Fanfani, J. Fischer, K. H. Shutta, P. Mandros, D. L. DeMeo, J. Quackenbush, and C. M. Lopes-Ramos. 2024a. Gene regulatory networks reveal sex difference in lung adenocarcinoma. Biol. Sex Differ. 15, 1 (Aug. 2024), 62.

59. E. Saha, V. Fanfani, P. Mandros, M. B. Guebila, J. Fischer, K. H. Shutta, D. L. DeMeo, C. M. L. Ramos, and J. Quackenbush. 2024b. Bayesian inference of sample-specific coexpression networks. Genome Research (2024), gr–279117.

60. R. Schulte-Sasse, S. Budach, D. Hnisz, and A. Marsico. 2021. Integration of multiomics data with graph convolutional networks to identify new cancer genes and their associated molecular mechanisms. Nature Machine Intelligence 3, 6 (June 2021), 513–526.

61. T. Schumacher, H. Wolf, M. Ritzert, F. Lemmerich, J. Bachmann, F. Frantzen, M. Klabunde, M. Grohe, and M. Strohmaier. 2020. The Effects of Randomness on the Stability of Node Embeddings. (May 2020). 2005.10039 [cs, stat].

62. D. Schwabe, S. Formichetti, J. P. Junker, M. Falcke, and N. Rajewsky. 2020. The transcriptome dynamics of single cells during the cell cycle. Mol Syst Biol 16, 11 (Nov. 2020), e9946.

63. P. Shannon, A. Markiel, O. Ozier, N. S. Baliga, J. T. Wang, D. Ramage, N. Amin, B. Schwikowski, and T. Ideker. 2003. Cytoscape: a software environment for integrated models of biomolecular interaction networks. Genome Res 13, 11 (Nov. 2003), 2498–2504.

64. K. H. Shutta, D. Weighill, R. Burkholz, M. Guebila, D. L. DeMeo, H. U. Zacharias, J. Quackenbush, and M. Altenbuchinger. 2022. DRAGON: Determining Regulatory Associations using Graphical models on multi-Omic Networks. Nucleic Acids Research 51, 3 (12 2022), e15.#x2013;e15. DOI:10.1093/nar/gkac1157 arXiv:https://academic.oup.com/nar/article-pdf/51/3/e15/49192710/gkac1157.pdf

65. A. Sibille, J.-L. Corhay, R. Louis, V. Ninane, G. Jerusalem, and B. Duysinx. 2022. Eosinophils and Lung Cancer: From Bench to Bedside. International Journal of Molecular Sciences 23, 9 (2022). DOI:http://dx.doi.org/10.3390/ijms23095066

66. D. Silverbush, S. Cristea, G. Yanovich-Arad, T. Geiger, N. Beerenwinkel, and R. Sharan. 2019. Simultaneous integration of multi-omics data improves the identification of cancer driver modules. Cell Syst. 8, 5 (May 2019), 456–466.e5.

67. P. Silveyra, N. Fuentes, and D. E. Rodriguez Bauza. 2021. Sex and Gender Differences in Lung Disease. Springer International Publishing, Cham, 227–258. DOI:10.1007/978-3-030-68748-9_14

68. D. A. Stroud, L. E. Formosa, X. W. Wijeyeratne, T. N. Nguyen, and M. T. Ryan. 2012. Gene knockout using transcription activator-like effector nucleases (TALENs) reveals that human NDUFA9 protein is essential for stabilizing the junction between membrane and matrix arms of complex I. J Biol Chem 288, 3 (Dec. 2012), 1685–1690.

69. A. Subramanian, P. Tamayo, V. K. Mootha, S. Mukherjee, B. L. Ebert, M. A. Gillette, A. Paulovich, S. L. Pomeroy, T. R. Golub, E. S. Lander, and J. P. Mesirov. 2005. Gene set enrichment analysis: A knowledge-based approach for interpreting genome-wide expression profiles. Proceedings of the National Academy of Sciences 102, 43 (Oct. 2005), 15545–15550.

70. D. Szklarczyk, A. L. Gable, K. C. Nastou, D. Lyon, R. Kirsch, S. Pyysalo, N. T. Doncheva, M. Legeay, T. Fang, P. Bork, L. J. Jensen, and C. von Mering. 2021. The STRING database in 2021: customizable protein-protein networks, and functional characterization of user-uploaded gene/measurement sets. Nucleic Acids Research (Database issue) 49 (2021).

71. S. J. Thomas, J. A. Snowden, M. P. Zeidler, and S. J. Danson. 2015. The role of JAK/STAT signalling in the pathogenesis, prognosis and treatment of solid tumours. British Journal of Cancer 113, 3 (July 2015), 365–371.

72. J. A. Tropp. 2012. User-friendly tail bounds for sums of random matrices. Foundations of computational mathematics 12 (2012), 389–434.

73. M. Udell and A. Townsend. 2019. Why are big data matrices approximately low rank? SIAM Journal on Mathematics of Data Science 1, 1 (2019), 144–160.

74. P. Veličković, G. Cucurull, A. Casanova, A. Romero, P. Liò, and Y. Bengio. 2017. Graph Attention Networks. In ICLR 2018. http://arxiv.org/abs/1710.10903

75. C. Wang, W. Rao, W. Guo, P. Wang, J. Liu, and X. Guan. 2022. Towards Understanding the Instability of Network Embedding. IEEE Transactions on Knowledge and Data Engineering 34, 2 (Feb. 2022), 927–941.

76. D. Weighill, M. Ben Guebila, K. Glass, J. Platig, J. J. Yeh, and J. Quackenbush. 2021. Gene targeting in disease networks. Front. Genet. 12 (April 2021), 649942.

77. C. Wilks, S. C. Zheng, F. Y. Chen, R. Charles, B. Solomon, J. P. Ling, E. L. Imada, D. Zhang, L. Joseph, J. T. Leek, A. E. Jaffe, A. Nellore, L. Collado-Torres, K. D. Hansen, and B. Langmead. 2021. recount3: summaries and queries for large-scale RNA-seq expression and splicing. Genome Biology 22, 1 (Nov. 2021), 323.

78. W. Yang, J. Soares, P. Greninger, E. J. Edelman, H. Lightfoot, S. Forbes, N. Bindal, D. Beare, J. A. Smith, I. R. Thompson, S. Ramaswamy, P. A. Futreal, D. A. Haber, M. R. Stratton, C. Benes, U. McDermott, and M. J. Garnett. 2012. Genomics of Drug Sensitivity in Cancer (GDSC): a resource for therapeutic biomarker discovery in cancer cells. Nucleic Acids Res 41, Database issue (Nov. 2012), D955–61.

79. B. Yu and K. Kumbier. 2020. Veridical data science. Proceedings of the National Academy of Sciences 117, 8 (2020), 3920–3929. DOI:10.1073/pnas.1901326117 arXiv:https://www.pnas.org/doi/pdf/10.1073/pnas.1901326117

80. C. Yu, A. M. Mannan, G. M. Yvone, K. N. Ross, Y.-L. Zhang, M. A. Marton, B. R. Taylor, A. Crenshaw, J. Z. Gould, P. Tamayo, B. A. Weir, A. Tsherniak, B. Wong, L. A. Garraway, A. F. Shamji, M. A. Palmer, M. A. Foley, W. Winckler, S. L. Schreiber, A. L. Kung, and T. R. Golub. 2016. High-throughput identification of genotype-specific cancer vulnerabilities in mixtures of barcoded tumor cell lines. Nature Biotechnology 34, 4 (April 2016), 419–423.

81. Y. Yu, T. Wang, and R. J. Samworth. 2015. A useful variant of the Davis–Kahan theorem for statisticians. Biometrika 102, 2 (2015), 315–323.

